# Multiplex-edited mice recapitulate woolly mammoth hair phenotypes

**DOI:** 10.1101/2025.03.03.641227

**Authors:** Rui Chen, Melissa L. Coquelin, Kanokwan Srirattana, Rafael Vilar Sampaio, Jacob Weston, Rakesh Ganji, Raphael A. Wilson, Jessie Beebe, Alba Ledesma, Jacob Sullivan, Yiren Qin, J. Chris Chao, James B. Papizan, Anthony Mastracci, Ketaki Bhide, Jeremy Abraham Mathews, Gregory Knox, Rorie Oglesby, Mitra Menon, Tom van der Valk, Austin Bow, Brandi L. Cantarel, Matt W. James, James Kehler, Love Dalén, Ben Lamm, George M. Church, Beth Shapiro, Michael E. Abrams

**Author notes:** These authors contributed equally to this work (Rui Chen, Melissa L. Coquelin).

## Abstract

The woolly mammoth (*Mammuthus primigenius*) possessed a thick woolly coat and other cold-adaptive traits that enabled survival in harsh arctic environments. Current de-extinction efforts focus on genetically modifying the closely related Asian elephant to express woolly mammoth traits. In this study, we establish a multiplex-edited mouse model with modifications in genes associated with hair morphology and lipid metabolism, enabling insights into traits involved in developing woolly hair textures. Our optimized workflows achieved high editing efficiencies and produced genetically modified mice with simultaneous editing of up to seven different genes. Selected modifications include loss-of-function mutations in *Fgf5, Tgm3*, and *Fam83g*, among others. The resulting mice display exaggerated hair phenotypes including curly, textured coats, and golden-brown hair. This study advances methods of rapid establishment of complex genetic models. These approaches inform de-extinction efforts and research involving the genetic basis of mammalian hair development.

## INTRODUCTION

One of the most prominent de-extinction efforts focuses on the woolly mammoth, *Mammuthus primigenius*, which disappeared following the extinction of the last surviving population on Wrangel Island approximately 4,000 years ago^1,2^. The mammoth de-extinction project aims to resurrect key mammoth traits such as woolly coats and other adaptations to cold habitats by genetically modifying mammoth-specific gene variants in the cells of the closely related Asian elephant, *Elephas maximus*. Genetic variants underlying phenotypes of interest are selected through comparative genomic analysis between the mammoth and elephant genomes^3^. Late-stage phases of the project involve somatic cell nuclear transfer of genetically modified Asian elephant cells, and subsequent surrogacy^4^. While active de-extinction efforts are underway with Asian elephant cells *in vitro*, phenotypic validation at the organism level faces significant challenges. The 22-month gestation period of elephants as well as their extended maturation timeline make rapid trait assessment impractical. Furthermore, ethical considerations regarding the experimental manipulation of elephants, an endangered species with complex social structures and high cognitive capabilities, necessitate alternative animal systems for proof-of-principle studies.

Mice are a well-defined model species with a 20-day gestation period, offering an ideal platform for refining methodologies and pipelines relevant for de-extinction. Specifically, workflows spanning targeted trait engineering, multiplex genome editing, high-efficiency animal generation, and non-invasive phenotypic assays can be rapidly implemented in mouse model systems. To this end, the CRISPR/Cas9 system has revolutionized the development of genetically modified mouse models^5,6^, and over the last decade, an increasingly versatile toolkit for precise genetic modifications has been developed^7,10^. Multiple established techniques enable generation of gene-edited mice, including injection of CRISPR/Cas9-modified murine embryonic stem cells (mESCs) into blastocysts to generate chimeric animals^11^, or direct zygote editing through CRISPR/Cas9 delivery^5,12,13^.

Recent advances in genetic technology have improved our ability to create complex genetic models in mice, but challenges remain in making this process faster and more efficient. Traditional methods that inject mESCs into blastocysts only produce partially edited first-generation animals, requiring extensive breeding programs to create fully modified mice. While directly editing zygotes speeds up production time, it often results in mosaic patterns where not all cells contain the desired modifications^14,15^. Scientists have recently reduced mosaicism by injecting edited stem cells into 8-cell stage embryos^16,17^, but this still requires the time-consuming process of creating fully modified stem cell lines with normal chromosomes. Improving these genetic engineering workflows is important not only for de-extinction projects but also benefits the broader scientific and industrial communities.

Here, we used a mouse model to 1) validate our multiplex genetic engineering pipelines; 2) demonstrate organism-level proof-of-principle trait engineering; and 3) produce mammoth-like genetic edits. We selected hair development as the primary trait of interest in this study. This rationale is multifactorial: hair offers a visual and diverse phenotype readout^18^, murine hair development is well-characterized^19^, and one of the most defining traits of the woolly mammoth is its dense coat^20^. Previous studies of mice have revealed mutations in genes that regulate distinct properties of hair, such as length (*Fgf5*^21^), pattern (*Tgm3*^22^, *Fam83g*^23^, *Fzd6*^24^, *Krt25*^25^, *Krt10*^26^, *Tgfa*^27^), and color (*Mc1r*^28^) (**Figure 1A**). Using a trait engineering approach, we generated multiplex-edited mice with combinations of these mutations to evaluate whether a woolly coat phenotype could be engineered. We also selected two editing targets, *Krt27* and *Fabp2*, that parallel mammoth-specific variants from comparative analyses between mammoth and elephant genomes. We used optimized direct zygote editing (Experiments A-C) and high-efficiency chimera approaches (Experiments D-F) (**Figure 1B, Table S1**), enabling rapid (4-6 week) generation of multiplex-edited mice with as many as eleven edits in seven different genes. These novel combinations of genetic modifications yielded mice with varied hair phenotypes including woolly coat textures.

**Figure 1.**
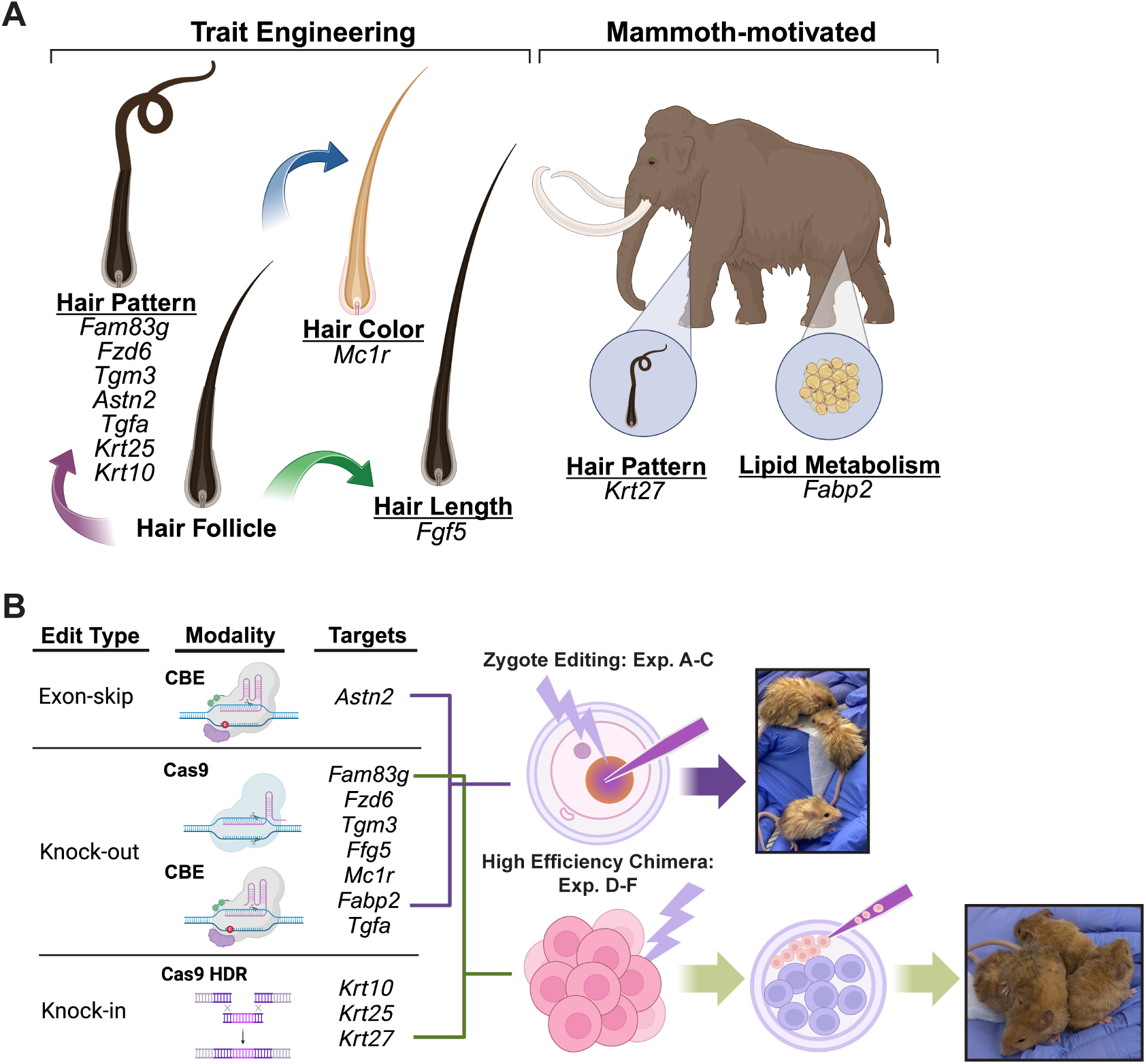
Overview of target traits, target genes, and editing workflows. Also see Table S1. **(A)** The target genes in this study fall into two categories: 1) woolly hair trait engineering based on existing literature, and 2) mammoth-motivated edits. **(B)** An illustration summarizing the editing modalities and two correlating animal generation workflows.

## RESULTS

### Rapid Generation of Woolly Mice via Multiplex Editing of Zygotes with CRISPR/Cas9

Electroporation of CRISPR/Cas9 ribonucleoprotein (RNP) complexes into mouse single-cell zygotes achieved efficient disruption of the six targeted genes: *Fgf5*, *Mc1r*, *Fam83g*, *Fzd6*, *Tgm3, and Fabp2* (**Figures 1B and 2A-B**). We carried out screening assays to prioritize sgRNAs based on three criteria: 1) high editing efficiency, 2) low predicted risk of off-target edits, and 3) targeting positions that either closely mimic previously reported mutations or that are in an early exon to facilitate full knock-out (KO).

**Figure 2.**
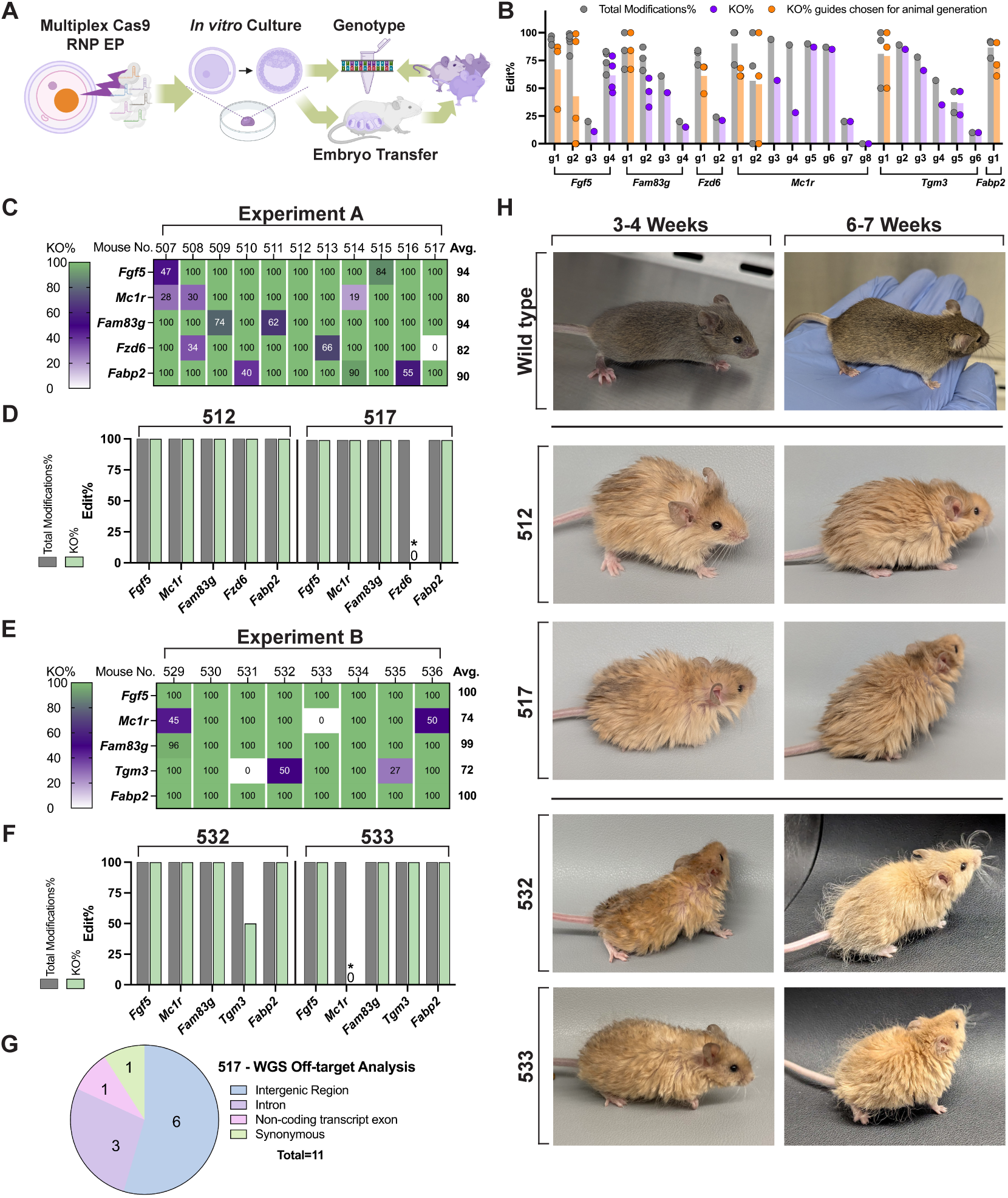
Experiments A and B: Gene editing in mice using CRISPR/Cas9 RNP zygote electroporation. **(A)** Workflow. We electroporated (EP) a pool of Cas9 RNPs with synthetic sgRNAs into wild-type mouse zygotes. We then cultured embryos into blastocyst stage *in vitro* and genotyped blastocysts to estimate editing efficiencies if needed. Next, we transferred blastocysts into surrogates for animal production experiments. Finally, we collected and genotyped ear punches from pups 21 days after birth. **(B)** Guide screening. We screened guides for editing efficiency in terminal experiments. For each guide, mean total modification efficiency (KO + unintended edits) is presented as a gray bar, KO efficiency as a purple bar for guides not selected for animal production experiments, and orange bar for guides selected for animal production experiments. Bars present average efficiencies from at least five embryos. Each circle represents an individual experiment that includes data from single embryos to pools of up to 18 embryos. **(C, E)** The KO% profiles of mice generated from Experiments A and B. Each column represents one mouse. “Mouse No.” is the name of the mouse used in this study. **(D, F)** The total modifications% and KO% from mice #512 and #517 of Experiment A and #532 and #533 of Experiment B. “*” highlights the two in-frame homozygous edits in #517 (a 9 bp deletion in *Fzd6*) and #533 (a 21 bp deletion in *Mc1r*). **(G)** Editing off-target analysis by whole genome sequencing on mouse #517. **(H)** Photos of mice #512 and #517 of Experiment A and #532 and #533 of Experiment B at 3-4 weeks old and 6-7 weeks old. The wild-type reference photos are from age-matched agouti B6SJL F2.

To assess editing efficiency for multiple sgRNAs per target gene, we electroporated thawed B6C3F1 × B6D2F1 zygotes with Cas9 RNP complexes. We then cultured embryos to the blastocyst stage, followed by collection for genotyping (**Figure 2A**). **Figure 2B** shows the editing efficiencies (% total modifications and % KO) of each sgRNA, which ranged from undetectable to 100% editing and highlights sgRNAs selected for subsequent animal production in orange. Binding sites of the selected sgRNA within each target gene are shown in **Figure S1A**.

We carried out two Cas9 editing animal production experiments using the selected sgRNAs. In Experiment A, we electroporated freshly collected B6SJLF2 zygotes with Cas9 RNPs and sgRNAs targeting *Fgf5*, *Mc1r*, *Fzd6, Fam83g*, *Fabp2*. Nearly half of the zygotes (63/134) developed to the blastocyst stage following electroporation (**Table S2**). We then transferred 60 blastocysts to three ICR pseudo-pregnant surrogates (**Figure 2A**). Eleven pups were born (18%; **Table S2**). Genotyping confirmed successful KO across all five target genes, with an average KO efficiency per gene across the 11 pups ranging from 80% for *Mc1r* to 94% for *Fam83g* (**Figure 2C**). Only modest mosaicism was detected: 54/55 of the possible target sites (5 targets x 11 mice) were 100% modified, and 42/55 had KO at 100%. The detailed genotypes of two representative mice, #512 and #517, are shown in **Figure 2D** and the remaining members of the litter are displayed in **Figure S1C**. All five target genes were fully disrupted in mouse #512, while mouse #517 had homozygous KOs for four target genes and a homozygous in-frame 9 bp deletion in *Fzd6*.

In Experiment B, we produced mice with simultaneous disruption (KO) of *Fgf5*, *Mc1r*, *Fam83g*, *Tgm3,* and *Fabp2* (**Figure 2E**). We electroporated zygotes as in Experiment A, resulting in a blastocyst formation rate of 56% (42/75; **Table S2**). Of these blastocysts, 28 were transferred to two ICR surrogates and eight pups were born, yielding a 29% birth rate (8/28; **Table S2**). Comparable levels of editing were achieved for the four gene targets in common with Experiment A. We detected little mosaicism, as 40/40 possible sites (8 mice x 5 targets) were fully modified, and only 6/40 had less than 95% KO. The unique target gene for this experiment, *Tgm3*, was edited at 100% in all eight mice, with 100% KO in five out of eight. The detailed genotypes of two mice, #532 and #533, are shown in **Figure 2F**, and the remaining members are shown in **Figure S1D**.

We performed short-read whole genome sequencing of mouse #517 to investigate the frequency of off-target edits. For the selected guides, computational analysis predicted 282 sites with three or fewer guide mismatches against the mouse genome where off-target edits were most likely to occur. We identified only 11 off-target edits in mouse #517 at these sites, none of which resulted in missense mutations (**Figure 2G**).

Experiments A and B produced mice with a variety of coat and eye colors determined by the status of *Mc1r* editing, and the genetic background of other coat/eye alleles in B6SJL F2 individuals (**Figures 2H** and **S1B-D**). Wild-type B6SJL F2 coat colors include agouti, black, white, light gray, and cream depending on which alleles for *Asip*, *Oca2*, and *Tyr* are inherited (**Figure S1B**)^29,30^. *Oca2^p/p^*mice have red eyes and light gray or cream coats, depending on which agouti (*Asip*) allele is present^31,32^, while *Tyr^c/c^* mice have red eyes and white coats^33^. Unedited mice with agouti or black coats have black eyes^34^. *Mc1r* regulates pheomelanin and eumelanin and our intended disruption of *Mc1r* should only alter black or agouti coats to gold coats in black-eyed animals^28,35^.

We successfully knocked out *Mc1r,* resulting in gold mice with black eyes. This phenotype is observed in mice #511, #512, #513, #516, #517, #532, #533, #534, and #535. Mice #509, #510, #515, and #530 were also homozygous knockouts at *Mc1r* but had white coats (**Figures S1C-D**), as expected in mice variants for albino coats^34^. Two mice with full editing of *Mc1r* did not show the expected coat color phenotype. Mouse #533 had a homozygous 21 bp in-frame deletion (**Figures 2F and 2H**), which was not considered a KO conventionally. However, mouse #533 still had a gold coat, indicating that the in-frame mutation nevertheless disrupted *Mc1r* function. Mouse #536 had a large deletion KO in one *Mc1r* allele, and also a 21 bp in-frame deletion in the other allele but had black hair (**Figure S1D**). The 21 bp *Mc1r* deletion in mouse #536 was, however, shifted by 9 bp compared to that of mouse #533. When we aligned both mutations to the wild-type MC1R protein (**Figure S1E**), we observed that the deletion in mouse #536 occurred near the second transmembrane domain of MC1R and spans the E92 residue, the loss of which has been associated previously with black coat color^36^.

Both Experiments A and B generated founder mice with distinctive hair phenotypes, including increased length, altered texture, and enhanced waviness compared to wild type (**Figures 2H** and **S1C-D**). These phenotypes were consistent with previous reports describing disruptions to the individual target genes. For instance, *Fgf5* KO mice exhibited increased guard hair length, probably due to disruption of *Fgf5-*delayed anagen-catagen transition^21^. We observed a “feathered” hair texture in our *Fam83g-*edited mice, similar to the *wooly* (*wly*) phenotype^23^. Mutations in *Tgm3* have previously been associated with changes to hair texture^22^, and Experiment B mice with homozygous editing of *Tgm3* also exhibited wavy coats. Deletion of *Fzd6* has been shown to alter hair follicle orientation^24^, and all mice with *Fzd6* knocked out displayed an exaggerated hair pattern phenotype. Given our shotgun-style multiplex strategy and the subsequently high gene editing efficiencies, it is difficult to assess the contribution of *Fzd6* disruption in the background of other gene knockouts. However, mice #517 (**Figures 2F and 2H**) and #513 (**Figure S1C**) had incomplete disruption of *Fzd6,* either as an in-frame mutation (#517) or mosaicism (#513) yet still presented with a woolly coat. These results suggest that *Fzd6* disruption did not significantly alter the macroscopic hair texture in our mouse model, and future experiments will be needed to fully characterize hair follicle orientation. Loss of *Fzd6* function did not compromise any exaggerated hair phenotype seen in Experiment A. Overall, the disruption of selected hair genes led to the successful generation of trait-engineered mice with woolly, gold coats.

Most mice from Experiment A and B had successfully engineered truncations of *Fabp2*, a mammoth-motivated gene candidate involved in fatty acid trafficking (**Figure 1A**). Mammoths have a fixed mutation in FABP2, resulting in an early stop codon and truncated protein^37^. In mice, KO of *Fabp2* leads to sex-dependent metabolic perturbations and differences in adiposity^38,39^, suggesting a possible role in cold adaptation. All but two of the 19 mice in Experiments A and B were homozygously edited for the desired mutation of *Fabp2* (**Figures 2C and 2E**). In our small sample size, average mouse body mass did not significantly differ between gene-edited and wild-type mice fed a standard chow diet.

### Rapid Generation of Woolly Mice via Multiplex Editing of Zygotes with a Fluorescence enriched Cytosine Base Editor Approach

In Experiment C, we used a cytosine base editor (CBE) to KO seven genes simultaneously: *Fgf5*, *Mc1r*, *Fam83g*, *Fzd6*, *Tgm3*, *Astn2*, and *Fabp2* (**Figures 1B and 3A**). CBE deaminates cytosine (C) to thymidine (T) within a defined targeting region^40^. CBE can therefore precisely install premature stop codons (*Fgf5*, *Mc1r*, *Fam83g*, *Fzd6*, *Tgm3*, and *Fabp2*)^41^ or mutate splice sites for creating gene truncations (*Astn2*)^42,43^ without eliciting double-stranded breaks^40^.

**Figure 3.**
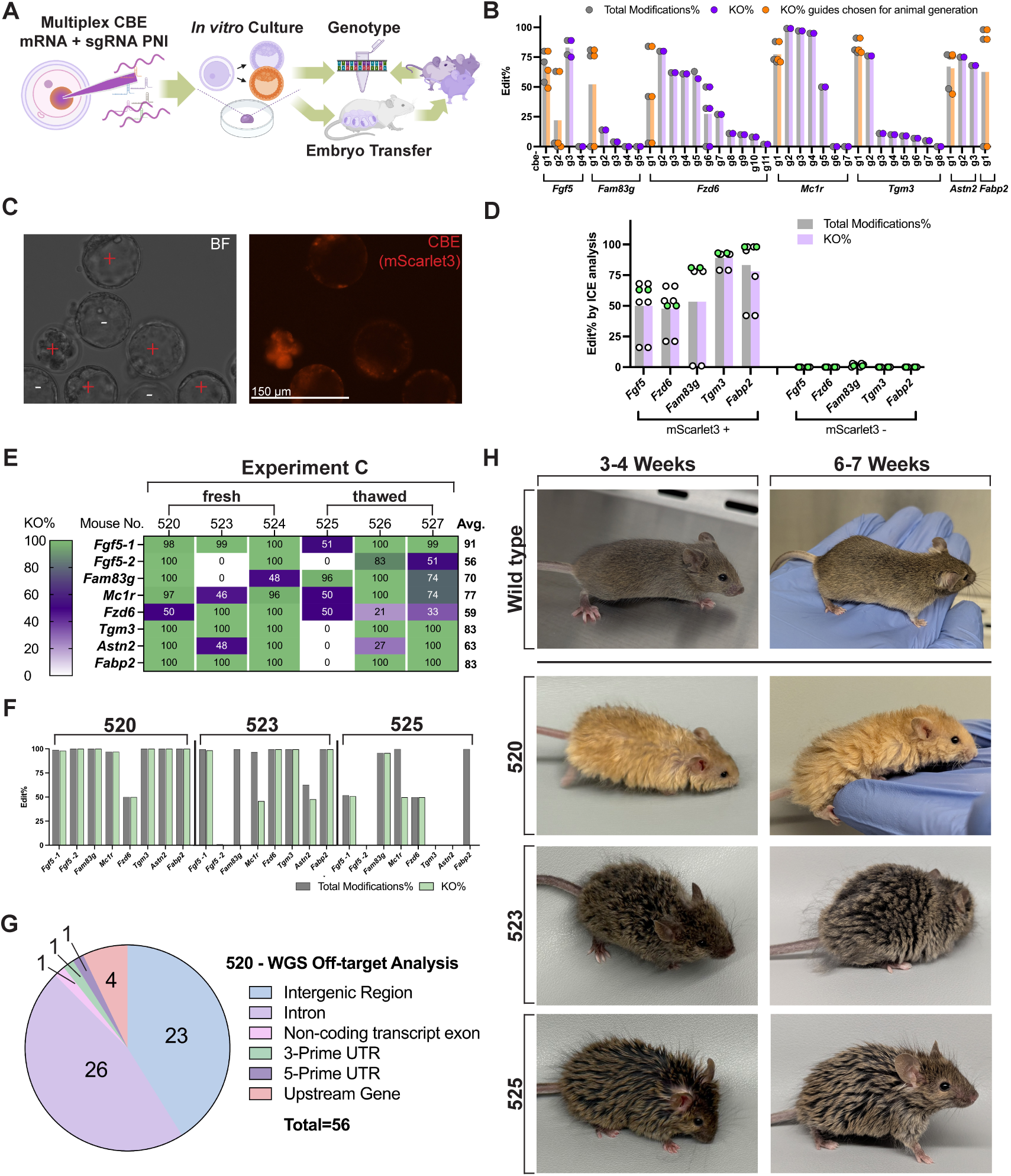
Experiment C: multiplex editing in mice using Cytosine Base Editor (CBE) PNI. **(A)** Workflow. We delivered a pool of CBE mRNA with sgRNAs into wild-type mouse zygotes by PNI. The mRNA transcript co-expresses CBE and a red fluorescent protein, mScarlet3. We cultured the embryos to the blastocyst stage *in vitro* and then evaluated mScarlet3 fluorescence. For screening experiments, we genotyped all blastocysts. For animal-producing experiments, we selected healthy blastocysts with bright mScarlet3 signals for transfer into surrogates. We collected ear punches for genotyping 21 days after the birth of the pups. **(B)** Guide screening. We screened guides for editing efficiency in terminal experiments. For each guide, mean total modification efficiency (KO + unintended edits) is presented as a gray bar, KO efficiency as a purple bar for guides not selected for animal production experiments, and an orange bar for guides selected for animal production experiments. Bars present average efficiencies measured from at least six embryos. Each circle represents an individual experiment, including data from single embryos to pools of up to 19 embryos. **(C)** Fluorescent images of blastocysts after PNI of CBE-P2A-mScarlet3 mRNA and sgRNAs. Right: red signal from mScarlet3; left: bright field (BF) showing all embryos. “+” highlights the red embryos, and “-” highlights the non-red embryos. **(D)** Mean total modifications% (grey bars) and KO% (purple bars) comparison between mScarlet3+ and mScarlet3-embryos. Bars include data from at least seven embryos. White circles represent data from single embryos; green circles represent data from 6-8 pooled embryos. **(E)** The KO% profiles of mice generated from the CBE animal production Experiment C. Each column represents one mouse. “Mouse No.” is the designated name of the mouse used in this study. **(F)** The total modification% and KO% from mouse #520, #523, and #525 from Experiment C. **(G)** Off-target analysis by whole genome sequencing on mouse #520. **(H)** Photos of mouse #520, #523, and #525, from Experiment C, taken at 3-4 weeks old and 6-7 weeks old. The wild-type reference photos are from age-matched agouti B6SJL F2; the same images as in Figure 2H.

To evaluate the efficiency of our CBE sgRNAs, we performed pronuclear injection (PNI) using thawed F2 hybrid zygotes with *in-vitro*-transcribed CBE mRNA and chemically synthesized sgRNA and collected embryos at the blastocyst stage for genotyping (**Figure 3A**). We found that cytosine deamination varied depending on the sgRNA and the target gene (**Figure 3B**). We prioritized sgRNAs for animal-production experiments (**Figure 3B** orange bars, **Figure S2A**) using the selection criteria described for Experiments A and B.

The co-expression of a red fluorescent protein, mScarlet3, via a P2A linkage was an effective reporter for CBE editing activity in zygotes (**Figures 3C-D and S2B-C**). We microinjected thawed F2 hybrid zygotes with CBE editor and multiplexed sgRNAs targeting *Fgf5*, *Fzd6*, *Fam83g*, *Tgm3*, and *Fabp2*. We manually sorted blastocysts into two groups based on the presence (+) or absence (-) of the red fluorescence signal. Genotyping analysis confirmed successful base editing in all red-fluorescent (+) embryos as well as high multiplexing in single embryos (**Figures 3D white dots, and S2B-C**). In contrast, embryos lacking red fluorescence (-) showed no detectable gene editing.

We leveraged the CBE reporter system to create mice edited at *Fgf5, Mc1r, Fam83g, Fzd6, Tgm3, Astn2*, and *Fabp2*. Fresh and thawed embryos exhibited comparable blastocyst formation rates: 70% (78/111) in fresh embryos and 71% (46/65) in thawed embryos (**Table S2**). Of these, the fluorescence rate of embryos was 65% (51/78) in fresh embryos and 35% (16/46) in thawed embryos. Five pups were obtained from the fresh zygote group after transferring 47 fluorescent embryos (11%; 5/47; **Table S2**), and three pups were born from the thawed group (33%; 3/9; **Table S2**). Every pup had at least heterozygous editing at more than one locus, and multiple pups were homozygous at 5 to 6 loci, with an average KO rate of 73% across six mice (**Figure 3E**). Average KO efficiencies per gene across all mice ranged from 59% for *Fzd6* to 91% for *Fgf5*. Mice #520 and #524 were edited at all seven genes. Mice #520 and #524 had homozygous KO for six genes and were heterozygous at a seventh, *Fzd6* and *Fam83g,* respectively (**Figure 3E**). Due to our high editing efficiency and the nuclease-free nature of CBE, total gene editing (intended edits + bystander edits + indels) often matched precision stop codon installation, as few unintended insertions or deletions were detected (**Figure 3F**, #520).

Multiplexed CBE editing in zygotes led to few off-target edits. Our WGS analysis of 1,869 computationally predicted off-target sites revealed 56 total off-target edits in mouse #520 (**Figure 3G**), none of which resulted in missense mutations. The higher number of predicted and observed off-target sites in Experiment C compared to Cas9 editing in Experiment A was as expected due to the less-stringent PAM of the SpRY-based^44^ editor that was employed, which provides flexibility in targeting but more promiscuous editing^45^.

Experiment C produced mice with similarly altered phenotypes as in Experiments A and B (**Figures 3H and S2D**), and both fresh (#520, #523, #524) and thawed (#525, #526, #527) embryos generated mice with comparable genotype to phenotype correlations. We observed that the addition of early stop codons in *Fgf5, Fam83g, Fzd6,* and *Tgm3* and skipping an exon in *Astn2* yielded an exaggerated hair phenotype (**Figures 3F and 3H**). Due to the lower editing efficiency of *Mc1r* in Experiment C compared to Experiments A and B, we obtained more agouti mice. Interestingly, while mouse #525 had only a single gene homozygously knocked out – *Fam83g –* this mouse still had a feathered coat phenotype (**Figure 3F and 3H**). Mouse #523 was not edited at *Fam83g* (**Figure 3F**), and did not display a feathered coat, but rather had a frizzy/curly hair phenotype (**Figure 3H**), which may be due to the additive effects of *Fgf5, Fzd6,* or *Tgm3* disruption.

### F0 Production of Woolly Mice with Near Full Contribution from Precision Edited mESCs

For Experiments D, E, and F, we established clonal mESCs modified with KO of *Fgf5*, *Mc1r*, *Fabp2, Tgfa*, *Fam83g, Fzd6,* and *Tgm3*, and knock-in (KI) of point mutations in the keratin family genes *Krt25*, *Krt27* and *Krt10* (**Figures 4A-B**), followed by high-yield chimera generation (**Figure 5A**). *Tgfa, Krt10, Krt25,* and *Krt27* were not genes modified in Experiments A-C. Tgfa is an EGFR ligand that plays a critical role in regulating hair development, and for which loss of function results in wavy hair textures^27,46^. We targeted an E172V substitution in *Krt10*, which has been associated with a “rough” coat texture, but normal hair length^26^. We separately targeted a L86Q substitution in *Krt25*, which has been linked to a wavy coat ^25^. On the other hand, mammoth-motivated gene candidate *Krt27* was selected from our genomic analysis ^37^, which revealed that this gene had fixed protein coding variants in mammoths compared to Asian elephants. Specifically, mammoths have a valine at amino acid position 191 of KRT27, while Asian elephants have a methionine. Mutations in *Krt27* have previously been linked to hair phenotypes in mice^47^. To replicate the mammoth variant at this site, we introduced a valine at the same position in mouse *Krt27*, where wild-type mice have isoleucine (**Figure S3A**). We designed sgRNAs for Cas9 or CBE to target each gene (**Figures 4C-D**) as well as single-stranded oligonucleotide donors (ssODN) to act as HDR templates for *Krt10*, *Krt25,* and *Krt27* (**Figure 4C**). We used optimally performing guides and ssODNs (**Figures 4C-D orange bars, and S3C-D**) for subsequent mESC editing experiments. For use in these experiments, we derived a B6129 mESC line from freshly obtained mouse zygotes, enabling a low passage number starting point (see **Methods**).

**Figure 4.**
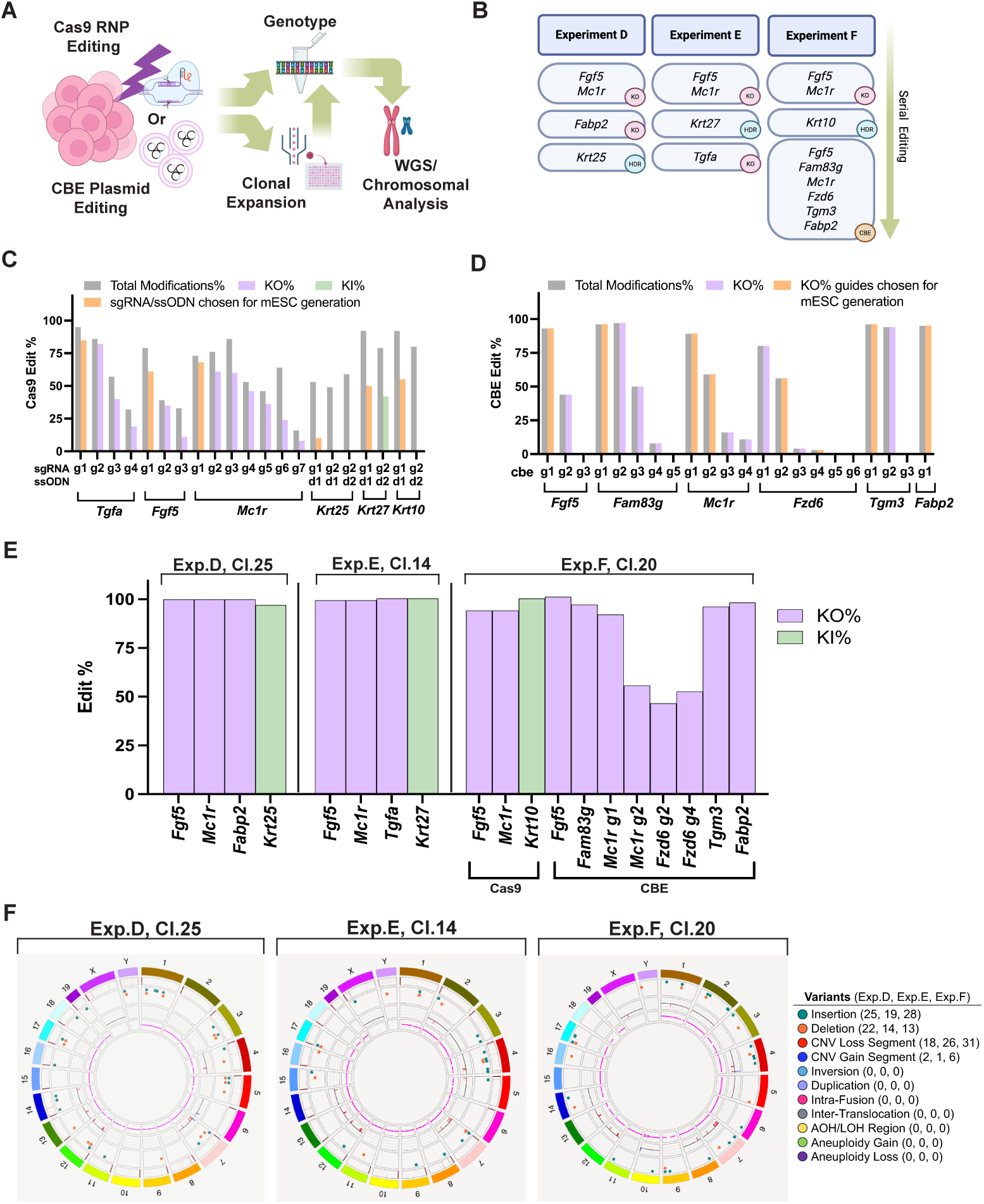
Experiment D, E, and F: generation of monoclonal, multiplex edited mouse embryonic stem cells using CRISPR/Cas9 RNP and CBE for woolly hair trait engineering. **(A)** Schematic for the generation of multiplex-edited monoclonal mESCs by CRISPR/Cas9 mediated KO or KI. We electroporated mESCs with Cas9 RNPs, sgRNAs and ssODN (HDR template) or used Lipofectamine to deliver CBE and guide RNA plasmids. We collected cells 48 hours post-transfection for genotyping by Sanger sequencing. We also single-cell-sorted the bulk population of transfected mESCs into 96-well plates containing iMEF feeders for clonal expansion, genotyping, and karyotyping. **(B)** Flowchart depicting the sequential editing strategy used to generate multiplex-edited monoclonal mESC lines. **(C, D)** Cas9 RNP guide, HDR ssODN, and CBE guide screening. Total modification efficiency is presented as a gray bar. KO or KI efficiency is presented as a purple or green bar, respectively, for designs not selected for animal production experiments, and an orange bar for designs selected for animal production experiments. Bars of each guide or each guide-ssODN combination includes data from one experiment from two thousand mESCs. **(E)** Bar graph showing KO% (purple) and KI% (green) for selected clonal lines generated in Experiments D, E, and F. We re-collected cells from laminin-coated plates to ensure no iMEF contamination during secondary genotyping by Sanger sequencing. Editing efficiencies were also confirmed by NGS. **(F)** Bionano Circos plot summarizing structural variations in Experiment D, E, and F monoclonal mESC lines compared to the unedited WT cell line. Starting from the outer rings, the Circos tracks depict 1. Cytoband, 2. mm39 DLE-1 SV mask, 3. Structural variants, 4. Copy number variants, and 5. Variant allele fraction segments. Translocations would be depicted as curved lines across the center of the plot, though none were detected in any sample. Numbers of variants are summarized on the right.

**Figure 5.**
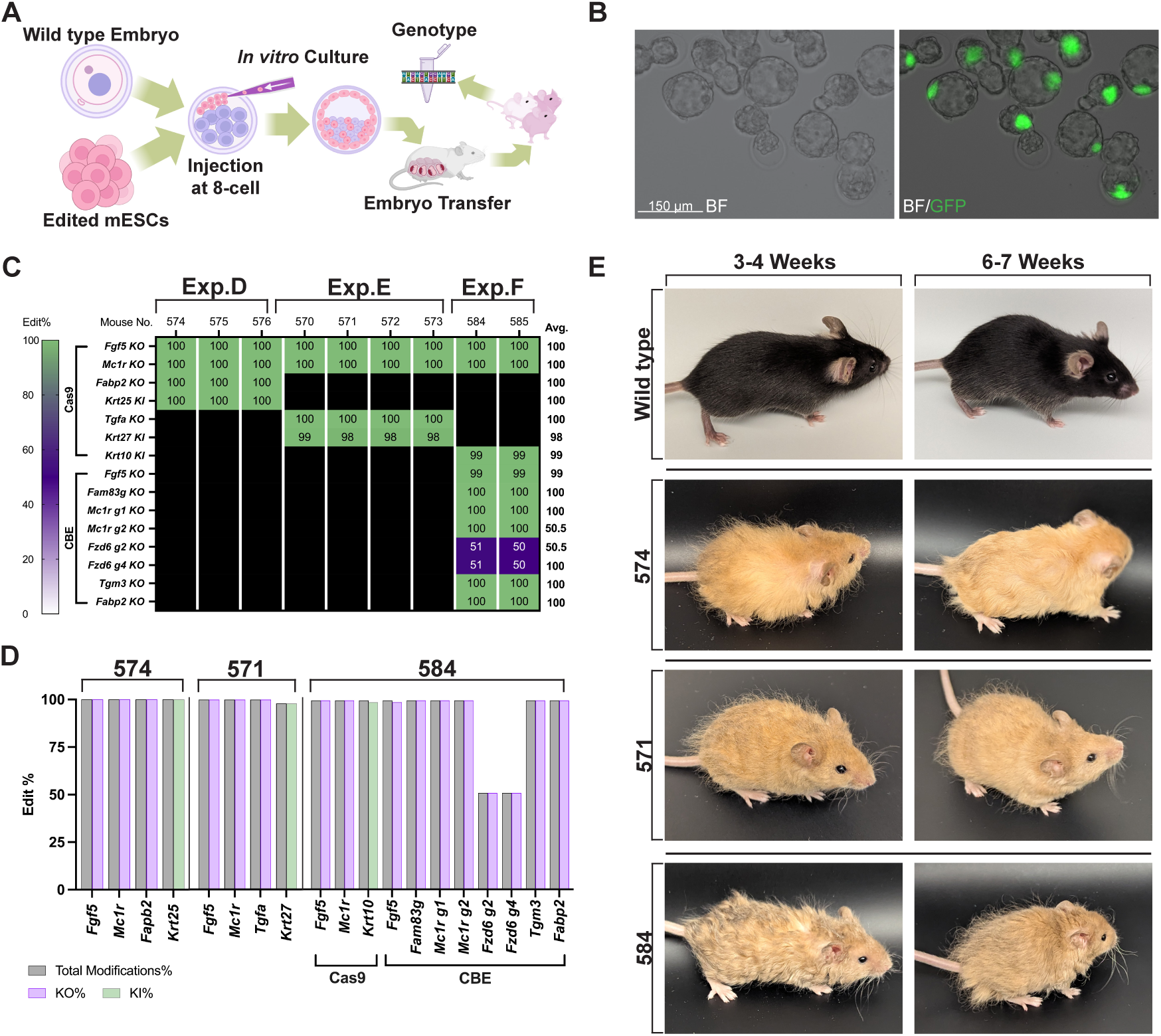
Experiment D, E, and F: generation of high yield woolly mouse chimeras using multiplex edited, monoclonal mouse embryonic stem cells. **(A)** Workflow for highly efficient production of edited mice within a single generation. We cultured edited monoclonal mouse ESCs in 2i medium and injected eight mESCs into an 8-cell stage zygote. We cultured the injected embryos in KSOM medium until the blastocyst stage and transferred the embryos to surrogates. Twenty-one days after birth, we collected ear punches for genotyping. **(B)** Representative image of blastocyst-stage embryos injected at the 8-cell stage with mESCs expressing GFP. Images taken at 20X. BF, bright field. **(C)** The KO% and KI% editing profile of mice generated from Experiment D, E, and F monoclonal cell lines. Each column is NGS data from n = 1 mouse. “Mouse No.” is the name of the mouse used in this study. **(D)** The total modifications%, KO%, and KI% from mice #574 (Experiment D), #571 (Experiment E), and #584 (Experiment F). **(E)** Photos of mice #574 (Experiment D), #571 (Experiment E), and #584 (Experiment F) at 3-4 weeks old and 6-7 weeks old. Wild-type reference photos are from an age-matched C57BL/6N mouse.

We developed edited clonal mESC lines via three serial transfections targeting three different panels of gene combinations among Experiment D, E, and F (**Figure 4B**), consisting of four, four, and eleven edits, respectively. Each of these experiments included serial editing by RNP delivery. Additionally, the final transfection for Experiment F involved plasmid-based editing by CBE and a pair of 4-guide RNA arrays, coupled with enrichment by the BEAR (Base Editor Activity Reporter) reporting system^48^ (**Figure S3B**). Post-transfection bulk cell populations exhibited edits at all desired loci with varying efficiencies (**Figure S3E**). Bulk cell populations were sorted to isolate single-cell clones, followed by genotype screening (**Figures 4A and S3F**). To identify fully edited clonal lines, we evaluated the editing status of 45 mESC clones for Experiment D, 48 clones for Experiment E, and 43 clones for Experiment F. To expedite the initial genotype screen, we retained mESCs on irradiated mouse embryonic fibroblasts (iMEF) feeder layers and genotyped them by Sanger sequencing (**Figures S4A**). The carry-over of iMEF wild-type DNA and the detection limitation of ICE analysis for Sanger results may explain the distribution of editing (non-50% multiples) obtained. We re-genotyped promising candidates after culture on feeder-free laminin coated plates. We proceeded with highly modified clones that exhibited robust proliferation and excellent colony morphology - Clone 25 for Experiment D, Clone 14 for Experiment E, and Clone 20 for Experiment F (**Figures 4E and S4B**).

Whole genome sequencing data from the fully edited mESCs clonal lines and wild-type mESCs revealed negligible off-target editing. As the edited clonal lines had the same genetic background (B6129), we were able to carry out an unbiased genome-wide off-target analysis by comparison to the unedited mESC line. We found limited high-impact off-targets, defined as splice modifying or protein coding variants. There were none in Experiment D (Clone 25), one in Experiment E (Clone 14), and none in Experiment F (Clone 20) (**Table S3**). Experiment E (Clone 14) had a single missense variant in the *Prkca* gene. The annotations for all variants found in this analysis are summarized in **Table S3**.

The candidate clonal lines selected for animal generation maintained normal chromosome numbers, as assessed by microscopy-based chromosome counting assays, PCR analysis, and Bionano optical genome mapping. To evaluate the three clonal lines for common mESC aneuploidies^49^, we performed Giemsa staining and subsequent chromosomal counting (**Figure S4C**), and found no detectable loss of chromosomes. PCR-based assessment of X and Y chromosomes showed retention of the Y chromosome (**Figure S4D**), the loss of which is a common mESC aneuploidy^49^. We also performed comprehensive structural genome analysis using Bionano (**Figure 4F**), which indicated that all 40 chromosomes were present in each of the cell lines with no apparent aneuploidy, translocations, or other genomic abnormalities. Copy number variants were minimal and often conserved between cell lines, including unedited B6129 mESCs, indicating that these variants were not caused by genetic editing.

The three edited mESC clones demonstrated robust high yield chimera formation, as each line produced mice with near full mESC contribution via injection into 8-cell stage embryos (**Figure 5A**). We first determined that mESCs localized to and formed the bulk of the inner cell mass by using a mESC line stably expressing eGFP (**Figures 5B and S3B**). Next, we injected mESC clone 25 (Experiment D), clone 14 (Experiment E), and clone 20 (Experiment F) (**Figure 4E**) into wild-type 8-cell embryos. We observed that 98% (147/150) of the 8-cell stage embryos injected reached the blastocyst stage after two days in culture, suggesting that *in vitro* embryo development was not negatively affected (**Table S2**). Between the three experiments, we generated nine mice all edited at the expected level at each target site (**Figure 5C**). The detailed genotypes of selected mice from Experiment D, E, and F are shown in **Figure 5D**, with the remaining mice shown in **Figure S5**. Given that all mice born were male and their editing profiles exactly matched that of the respective donor mESCs (**Figure 4E**), we infer that these mice were nearly 100% ESC-derived.

Mice from Experiments D, E, and F had a gold coat, long guard hairs, and distinctive woolly hair textures (**Figures 5E and S5**). Mice from Experiment D (*Fgf5, Mc1r, Fabp2* KO; *Krt25* KI) (**Figure 5E**, #574) had a rough, wiry coat with disoriented hair. This wiriness diminished after six weeks of age and became more “Plush” in texture, as reported previously for this *Krt25* mutation ^25^. Mice produced in Experiment E (*Fgf5, Mc1r, Tgfa* KO; *Krt27* KI) (**Figure 5E**, #571) had long, disordered guard hairs with a rough texture, as well as wavy hairs that remained closer in place to the body. Mice of Experiment F (*Fgf5*, *Mc1r*, *Fzd6*, *Fam83g*, *Tgm3*, *Fabp2* KO; *Krt10* KI) (**Figure 5E**, #584) had feathered, curly coats like mice from Experiment B or C, likely due to the shared targets. The phenotypes of mice produced within each experiment were strikingly similar to each other (**Figures 5E and S5A-C**), as expected due to the nearly full genetic contribution from their respective clonal mESC lines. Overall, this workflow produced nine founder F0 mice with near-full contribution from donor mESCs, with up to 11 edits across 7 genes and profound engineered woolly traits.

### Successful Germline Transmission by Both Zygote-edited and mESC-derived F0 woolly mice

We demonstrated successful germline transmission (**Figure 6**) produced by either zygote editing (Experiment A-B; **Figure 2**) or mESC-based high efficiency chimera creation (Experiment D-F, **Figures 4-5**). The F1 litters were healthy and had the expected genotype (**Figure 6A-D**) and phenotype (**Figure 6E**), corresponding to each dam/sire cross.

**Figure 6.**
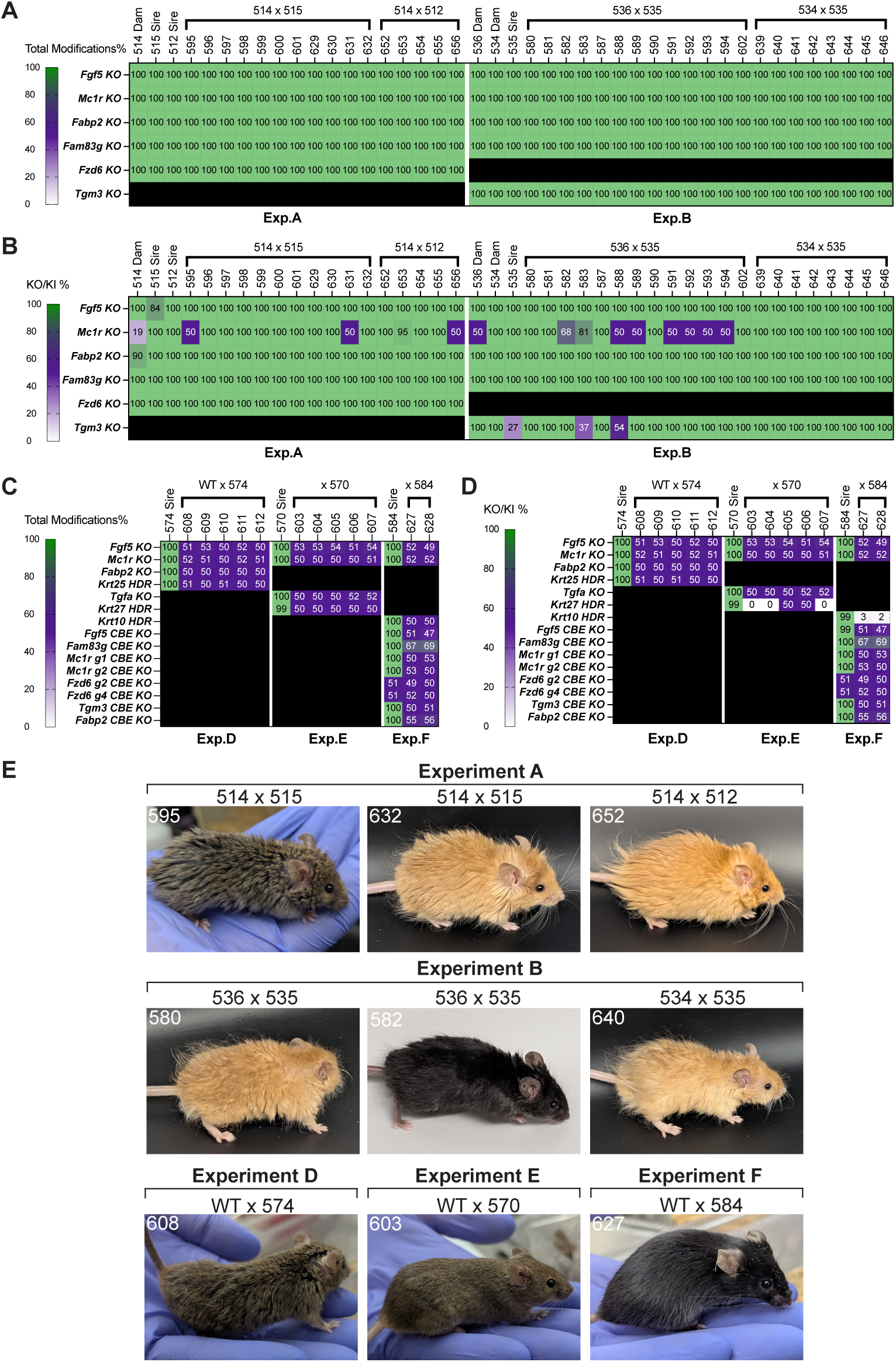
Germline editing is achieved by both zygote-edited and mESC-derived F0 woolly mice. (A, **C)** Total modification% or **(B, D)** desired edit (KI/KO) % of the dam, sire and their progeny from experiment A, B, D, E and F. #603, #604, #607, #627 and #628 inherited a large deletion on their HDR target, showing no KI. **(E)** Representative photos of F1 progeny from the breeding schemes of Experiment A and B (7-9 weeks) and Experiment D, E, and F (12-16 weeks).

To determine whether founder mice produced via zygote editing had germline gene modification, we set up two crosses for mice from Experiment A and two crosses for mice from Experiment B. We mated mouse #514 (dam) to either mouse #515 or #512 (sire). Dam #514 had ∼20% mosaic *Mc1r* KO, as determined by genotyping and observed by coat color pattern. All progeny from #514x#515 or #514x#512 were fully modified at each gene (**Figure 6A**). However, mice #656 (#514x#512 progeny), #595, and #631 (#514x515 progeny) had heterozygous KO of *Mc1r* (**Figure 6B**). Consistent with this genotyping data, all progeny from Experiment A founders had long woolly coat phenotypes, and all progeny but #595, #631, and #656 had golden coat color (**Figure 6E**, #632 and #652, etc). Only progeny heterozygous for *Mc1r* had agouti color (**Figure 6E**, #595, etc). Given these data, it is likely that dam #514 had higher *Mc1r* KO mosaicism in gamete tissue compared to global editing. For Experiment B, we mated mouse #535 (sire) to either mouse #534 or #536 (dam). Sire #535 had partial KO of *Tgm3*, dam #536 had partial *Mc1r* KO, as well as an allele with a unique in-frame truncation (**Figures 6A-B and S1E**), and dam #534 had homozygous KO for all targets. Progeny from the #534x#535 cross had 100% KO of each target gene (**Figure 6B**), and golden woolly coats (**Figure 6E**, #640, etc). When dam #536 was crossed to sire #535, all progeny were 100% modified across the five target genes, with expected alleles contributed from the dam and sire (**Figures 6A-B**). For example, the 21 bp in-frame deletion of *Mc1r* from the dam (#536) (**Figure S1E**) was inherited by half the progeny (8/14, **Figure 6B**, #582-3, #588-9, #591-4). The coats of pups with these genotypes were black (**Figure 6E**, #582, etc), further supporting the hypothesis that this *Mc1r* truncation spanning the E92 residue generates black coats in a dominant manner over agouti coats. The other progeny (**Figure 6E**, #580, etc.) showed the signature *Mcr1* KO-mediated gold coats, due to the frame-shifted *Mc1r* alleles from founders #536 and #535. Demonstrated by F0 breeding, our high efficiency zygote editing pipeline rapidly generated germ-line homozygous 5-plex gene knockout mouse models from a single generation.

Male F0 mice produced via mESC injection into 8-cell embryos (Experiment D, E and F) mated to wild-type females produced F1 progeny (12/12 pups) that were heterozygous for the edits from the respective sires (**Figure 6C**). Since *Mc1r* was heterozygous, the F1 progeny had either agouti (**Figure 6E**, #608, #603, etc) or black coats (**Figure 6E**, #627, etc), depending on the coat color genetics of the wild-type dam. Specifically, mice #574 and #570 were bred to agouti B6SJLF2 dams and #584 was bred to a black B6SJLF2 dam. All progeny from the WTx#574 cross (Experiment D, 5/5) had a rough coat phenotype (**Figure 6E**, #608, etc), while progeny from WTx#570 (Experiment E, 5/5) or WTx#584 (Experiment F, 2/2) matings displayed no overt coat phenotype (**Figure 6E**, #603, #627, etc). All Experiment D F1 progeny had the expected heterozygous KO or KI at all four targets (**Figure 6D**).

Some pups sired from mice of Experient E (sire #570) and F (sire #584) did not have the exact HDR-mediated knock-in allele expected from the respective sire (**Figure 6D**). To investigate, we re-examined the *Krt10* and *Krt27* loci in the DNA samples from these sires and their progeny and found a large 400-500bp deletion. This deletion was not detected by our NGS genotyping strategy, as we employed a 300 bp amplicon spanning the target site of interest (**Figure S6**). Our expanded genotyping data showed that sires #570 and #584 were hemizygous for the precision edit in *Krt27* (#570) or *Krt10* (#584), with a deletion in the alternate allele abolishing the primer binding site and therefore preventing amplification. Thus, progeny inherited either the large deletion (**Figure 6D**, #603, #604, #607, #627, #628) or the precision edit (**Figure 6D**, #605, #606). Nevertheless, our mESC-based high-efficiency chimera pipeline achieved complete germline penetration, correlating to the editing status of the sire.

## DISCUSSION

We used several genome editing approaches to generate mice with combinatorial knock-outs or knock-ins in eleven genes involved in hair development and lipid metabolism (**Figures 1A-B**). The target genes were selected based on hair-associated mutations previously observed in mice, and mammoth-specific alleles identified through comparative genomic analysis. Our genetically modified mice have hair phenotypes including wiry coats, wavy hair, altered hair color, and woolly coats (**Figures 2H, 3H, 5E, S1C, S1D, S2D, and S5A-C**). Our mouse models provide validated animal production pipelines and demonstrated proof-of-concept trait engineering relevant to woolly mammoth de-extinction.

Our optimized protocols and workflows enabled the rapid and efficient production of multiplexed mouse models (**Figure 1B**). We found extensive guide screening essential to maintaining cost-effectiveness and minimizing zygote collections when generating genetically modified animals. Direct zygote editing with CRISPR/Cas9 enabled simultaneous KO of up to five genes with editing efficiencies exceeding 90% (**Figures 2C and 2E**). Similarly, CBE facilitated precise editing of eight target sites from seven genes simultaneously, while the fluorescent protein expression served as an effective reporter of editing success (**Figures 3C-E**). Importantly, high efficiency zygote editing resulted in germline modification, rapidly enabling F0 crosses to generate homozygously edited F1 offspring (**Figure 6A-B**). Our last strategy involved a two-step process: first mESCs were edited *in vitro*, and fully modified clones were identified; these clones were then injected into 8-cell embryos yielding chimera pups (**Figures 4A and 5A**). This approach yielded mice derived almost entirely from mESCs (**Figure 5C**), dramatically reducing the time needed to produce fully engineered mice and allowing F0 characterization of phenotypes^12,13,17,50,51^. Using a combination of RNP Cas9 KO, HDR, and base editing to genetically modify mESCs, we produced animals with up to eleven edits across seven target genes. Further, our high efficiency chimera method produced F0 animals with fully modified germ cells, and all edits were successfully inherited by every F1 progeny (**Figure 6C-D**).

Our mouse models provide evidence supporting the role of several spontaneous or induced mutations in genes involved in hair development, including *Fam83g*, *Tgm3*, and *Krt25*. The previously described “wooly” (*wly*) mutation arose spontaneously in a Jackson Laboratory colony. This phenotype is thought to be caused by altered splicing that leads to a frameshift mutation in exon 4 of *Fam83g*, although the researchers did not confirm this hypothesis by targeted gene knock-out^23^. Our *Fam83g* sgRNA used in Experiment A produces a similarly truncated protein by targeting the first few nucleotides of exon 3. Our study provides the first genetic validation that truncation of *Fam83g* alters hair texture. In *Tgm3,* multiple recessive wellhaarig (*we*) mutations are associated with wavy coats and curly vibrissae^22^. The *we^4j^* variant has been fine-mapped to a 7 bp deletion in *Tgm3* that leads to a truncation after residue 512^22^. To our knowledge, the *we^4j^* mutation has not been recapitulated by targeted CRISPR/Cas9 knockout. Our *Tgm3* sgRNA (Experiment B) targets within 40 bp downstream of *we^4j^*, leading to a similar frameshift, supplying evidence that CRISPR/Cas9 mediated *Tgm3* truncation results in a wavy coat phenotype. The “Plush” mutation has previously been mapped to the *Krt25* gene in N-ethyl-N-nitrosourea (ENU)-induced mutant mice^25^. Mice from Experiment D had homozygous KI of L86Q in *Krt25*, precisely replicating the induced mutation. These mice also exhibited an initial wavy coat, followed by a more plush-like coat with aging (**Figure 5E**, #574, and **S5A**), consistent with previous observations^25^, and thus provides evidence implicating the phenotypic effect of this substitution in *Krt25*. Another variant in *Krt25*, “Rex”, has been reported to result in curly coat phenotype in both heterozygotes and homozygotes^52^. Our F1 progeny from Experiment D also possessed a ruffled coat texture (**Figure 6E**, #608), probably from the dominant effect of heterozygous *Krt25* mutation.

The multiplexed editing strategies and mouse models established here provide a foundation for evaluating complex combinations of genetic modifications leading to woolly hair phenotypes. Findings from these experiments will continue to inform prioritization of genetic modifications for elephant cell engineering and mammoth de-extinction. Of course, while mouse models provide an experimentally tractable system for evaluating the functional consequences of genetic modifications, hair follicle biology and physiological parameters of mice probably differ from those of proboscideans, as they differ from those of humans^19^. Additionally, the mammoth cold-adaptive phenotype would have involved complex physiological modifications beyond hair morphology, including changes in adipose distribution, vascular architecture, and metabolic thermoregulation. Nevertheless, the technical advances in multiplexed editing demonstrated here have implications for both basic research and therapeutic applications requiring precise orchestration of multiple genetic modifications.

## RESOURCE AVAILABILITY

### Lead Contact

Further information and requests for resources and reagents should be directed to and will be fulfilled by the lead contacts, Michael E. Abrams and Beth Shapiro.

### Materials availability

All unique/stable reagents generated in this study are available from the lead contact with a completed materials transfer agreement.

### Data and code availability

Any additional information required to reanalyze the data reported in this paper is available from the lead contact upon request.

## ACKNOWLEDGMENTS

The authors would like to thank Dr. Robert Hammer, John Ritter and the Transgenic Core at UTSW for their insight and support providing custom mouse model services. We would also like to thank Katie SoRelle, Faith McCorkle, and Lauren Pandolfo of the Colossal animal care team, and the staff at the Animal Resource Center at UTSW for their excellent care. We thank Dr. Eriona Hysolli for assistance with this work. Finally, we would like to thank Dr. Daniel Rokhsar, Dr. Andrew Pask, and Dr. Jun Wu for reviewing this work and for helpful comments on this manuscript. This work was supported by Colossal Biosciences.

## AUTHOR CONTRIBUTIONS

Conceived and designed the experiments, performed experiments, carried out data analysis: R.C., M.L.C., K.S., R.V.S., R.A.W., R.G., J.W., A.L., J.B., J.S., A.B., and M.E.A; Cell culture and molecular biology experiments: R.C., M.L.C., R.A.W., R.G., J.W., J.B., J.S., G.K., and R.O.; Embryology experiments: K.S., R.V.S., A.L., and Y.Q.; Animal work: R.A.W., K.S., J.K., and M.W.J.; Bionano data acquisition and analysis: A.M.IV; NGS data acquisition: J.C.C., J.B.P., and A.M.IV; Bioinformatics analysis: K.B., J.A.M., M.M., J.W., and B.L.C.; Writing: R.C., M.L.C., K.S., R.V.S., R.A.W., R.G., J.W., A.L., J.B., J.S., B.S., and M.E.A.; Project supervision: B.C., M.J., J.K., B.L., G.M.C., B.S., and M.E.A.; Project conceptualization: B.L., G.M.C., B.S., L.D., T.v.d.V., and M.E.A.; Funding acquisition: B.L.; All authors were involved in reviewing and revising the manuscript.

## SUPPLEMENTAL INFORMATION

**Document S1. Figures S1–S6 and Table S1-6** (Included in the main PDF)

## MATERIALS AND METHODS

### Animal housing and strains

Animal work described in this manuscript was approved and conducted under the oversight of the UT Southwestern Institutional Animal Care and Use Committee (IACUC) Dallas, TX, USA. All mice were housed in standard cages within a specific pathogen-free facility, maintained under a 12-hour light/dark cycle.

We used embryos from B6SJLF1 (The Jackson Laboratory, ME, USA) mice for zygote editing (Experiment A-C) and C57BL/6N (Charles River Laboratories, MA, USA) or B6SJLF1 mice for chimera generation (Experiment D-F). ICR (Inotiv, IN, USA) mice served as pseudopregnant recipients. We obtained frozen B6C3F1 × B6D2F1 embryos from Embryotech (MA, USA).

### Pronuclear embryo production

Female B6SJLF1 mice (8–12 weeks old) were superovulated by administering 7.5 IU of pregnant mare serum gonadotropin (PMSG; Prospec, Rehovot, Israel), followed 48 hours later by 7.5 IU of human chorionic gonadotropin (hCG; Sigma-Aldrich, MO, USA). These females were then mated with male B6SJLF1 mice and fertilized embryos were collected from the oviducts 20 hours after the hCG injection. We maintained the harvested embryos in KSOM medium (EmbryoMax^®^ Advanced KSOM Embryo Medium, Sigma-Aldrich, MO, USA). We obtained fertilized oocytes from C57BL/6N mice following the superovulation and mating protocol described above, except with 6.0 IU of PMSG followed by 6.0 IU hCG.

We thawed frozen B6C3F1 × B6D2F1 zygotes (Embryotech, MA, USA) at room temperature for 2 minutes, followed by washing and incubation in KSOM medium for 10 minutes at room temperature. We then cultured the zygotes in KSOM medium under mineral oil (Sigma-Aldrich, MO, USA) at 37°C in an atmosphere of 5% CO₂, 5% O₂, and 90% N₂ for one hour before proceeding with further experiments.

### Zygote electroporation and pronuclear injection

We prepared RNP complexes for electroporation by preparing a mixture consisting of 1 µM recombinant *S. pyogenes* Cas9 nuclease (Alt-R^®^ S.p. HiFi Cas9 Nuclease V3, IDT, IA, USA), 1 µM of each sgRNA (Synthego, CA, USA or IDT, IA, USA), and 4 µM Alt-R® Cas9 Electroporation Enhancer (IDT, IA, USA) in OPTI-MEM^®^ medium (Gibco, NY, USA). We placed zygotes (30–50) into a 0.1 cm-gap electroporation cuvette (Bio-Rad, CA, USA) containing 10 µL of RNP complexes and performed electroporation at room temperature using an ECM 2001 Electroporation System (BTX, MA, USA) with the following parameters: 30 V direct current, 3 ms pulse duration, and 5 pulses.

For pronuclear injection, we prepared a solution containing 200 ng/µL CBE mRNA and 25 ng/µL of each sgRNA (Synthego, CA, USA or IDT, IA, USA) using nuclease-free water (Synthego, CA, USA). We injected this solution into the pronuclei of zygotes placed in droplets of KSOM medium under mineral oil. We performed microinjections using a FemtoJet 4i microinjector (Eppendorf, Hamburg, Germany) and a Nikon ECLIPSE Ti2 inverted microscope (Nikon, Tokyo, Japan) equipped with micromanipulators (Narishige, Ibaraki, Japan).

After electroporation and pronuclear injection, we cultured embryos in KSOM medium under mineral oil at 37°C in an atmosphere of 5% CO₂, 5% O₂, and 90% N₂ for 4 days. We then transferred blastocyst-stage embryos into the uteri of pseudopregnant ICR mice at 2.5 days post coitum.

### Fluorescent imaging

We imaged fluorescent blastocysts injected with CBE-mScarlet3 mRNA or injected with mESC expressing GFP using the Invitrogen EVOS M7000 Imaging system (Thermo Fisher Scientific, MA, USA) equipped with a 20X objective lens and RFP and GFP LED cubes.

### Murine embryonic stem cell (mESC) derivation

We derived B6129 mESCs from hybrid F1-fertilized embryos generated by crossing C57BL/6J females with 129/S4 males. We collected one-cell zygotes from plugged females following treatment with PMSG (Prospec, Rehovot, Israel) and hCG (Sigma-Aldrich, MO, USA), which we then cultured in advanced KSOM medium (Sigma-Aldrich, MO, USA) until reaching the blastocyst stage at four days post-fertilization. We removed the zona pellucida from the blastocysts via brief exposure to Tyrode’s acidic solution (Sigma-Aldrich, MO, USA). Zona-free blastocysts were collected using a mouth pipette, washed three times in KSOM medium, and individually plated in 48-well plates containing inactivated mouse embryonic fibroblasts (iMEFs, Thermo Fisher Scientific, MA, USA) and ESC derivation medium. The ESC derivation medium consisted of Knockout Dulbecco’s Modified Eagle Medium (KO-DMEM, Gibco, MA, USA) supplemented with 20% Knockout Serum Replacement (KOSR, Gibco, MA, USA), recombinant mouse leukemia inhibitory factor (LIF, Sigma-Aldrich, MO, USA), L-glutamine (Sigma-Aldrich, MO, USA), penicillin/streptomycin (Sigma-Aldrich, MO, USA), non-essential amino acids (Sigma-Aldrich, MO, USA), and β-mercaptoethanol (Sigma-Aldrich, MO, USA). After 7–8 days of culture, blastocyst outgrowths were enzymatically dissociated using 0.025% trypsin solution (Sigma-Aldrich, MO, USA) for 10 minutes. We neutralized the trypsinization reaction with ESC culture medium and the cells were then seeded onto 24-well plates pre-coated with iMEFs. We subsequently expanded several ESC colonies to 6-well plates on iMEFs for further use. We characterized ESC clonal lines for genomic stability, and selected a karyotypical male line at passage 2 for editing and animal production in this study.

### mESC Culture

We maintained B6129 mESCs on DR4 iMEFs at 37°C with 5% CO_2_ in culture medium consisting of KO-DMEM (Gibco, MA, USA) supplemented with 15% KOSR (Gibco, MA, USA), 1% GlutaMAX (Gibco, MA, USA), 1% MEM Non-Essential Amino Acids (Gibco, MA, USA), 100 µM 2-Mercaptoethanol (Gibco, MA, USA), 1000 U/mL LIF (Sigma-Aldrich, MO, USA), and 1% penicillin/streptomycin (Gibco, MA, USA), changing the growth medium daily. We used collagenase (Gibco, MA, USA) to gently lift ESCs from the iMEF feeder layer, followed by a brief incubation in diluted Accumax (Thermo Fisher Scientific, MA, CA) to generate a single cell suspension for passaging and electroporation.

### Transfection of mESC cell lines

We formed RNP complexes by combining 3200 ng of *S. pyogenes* Cas9 nuclease (Alt-R® S.p. HiFi Cas9 Nuclease V3, IDT, IA, US), 400 pmol of sgRNA (Synthego, CA, USA or IDT, IA, USA), and 5 µL of Alt-R® Cas9 Electroporation Enhancer (IDT, IA, USA). For RNPs targeting HDR, we mixed an additional 400 pmol of single-stranded DNA oligos (IDT, IA, USA) and 5 µL of Alt-R® Cas9 Electroporation Enhancer (IDT, IA, USA) with Cas9 and sgRNA. Complexes were allowed to form for 10-20 minutes at room temperature.

For Cas9 editing, we electroporated RNP into 500K to 1M mESCs with the Lonza P3 Primary Cell 4D-Nucleofector X Kit L (Lonza, Basel, Switzerland) using program CG104, according to the manufacturer’s protocol. Post electroporation, we seeded mESCs back on DR4 iMEFs in growth medium supplemented with Y-27632 2HCL (Selleckem, TX, USA). For HDR experiments, we added 1 µM of Alt-R®HDR enhancer (IDT, IA, USA) to the medium.

For CBE editing, we transfected plasmids into mESCs with Lipofectamine 2000 (Invitrogen, MA, USA) according to the manufacturer’s protocol. Briefly, we combined 4000 ng (1000 ng per plasmid) of CBE editor, BEAR, and two multiguide cassette plasmids with Lipofectamine 2000 in OptiMEM reduced serum medium (Gibco, MA, USA) at a wt/vol ratio of 1/3. Following DNA-lipofectamine complex formation, we added the complex to 500K to 1M mESCs in suspension and incubated at 37°C with 5% CO_2_ for 30 minutes. Cells were then seeded back onto DR4 iMEFs following quick centrifugation to remove the DNA-lipofectamine complexes.

### Clonal isolation of edited mESC lines

After three serial rounds of electroporation as described above, we single-cell sorted mESCs directly onto a 96-well plate pre-coated with iMEFs using the SONY SH800S sorter equipped with a 100 µM chip (SONY, CA, USA). For the CBE editing experiment, only BEAR (GFP) and editor (mScarlet3) double positive cells were selected for clonal outgrowth. We then expanded mESC colonies to a 24-well plate pre-coated with DR4 iMEFs until approaching confluency. Ten percent of the mESCs were collected for Sanger sequencing. The remaining ESCs were either cryopreserved in 90% KOSR/10% DMSO or seeded on laminin-coated plates (Laminin-511, BioLamina, Sundyberg, Sweden) for downstream analysis. Selected mESC clones for animal generation were thawed and expanded ten days before embryonic injection.

### Detection of Y chromosome by PCR

We performed PCR on mESC clonal expansions to detect loss of the Y chromosome following established primer designs^53^. We used a chromosomally normal male B6129 ESC line with an intact Y chromosome as a positive control and a male B6129 ESC line with known Y chromosome loss as a negative control. We performed a 15 µl PCR using KOD ONE (Sigma-Aldrich, MO, USA) as follows: 38-40 cycles of 98°C for 15 seconds, 60°C for 5 seconds, and 68°C for 5 seconds. We ran 10 µl of the resulting product on a 2% agarose gel at 220 volts for 30 minutes, and obtained a gel image using the iBright. (Thermo Fisher Scientific, MA, USA). Assuming no loss of the Y, the result should be a Y-chromosome specific amplicon (280 bp) and two amplicons specific to the X chromosome (685 bp and 480 bp).

### Chromosome counting assay

We performed chromosome counting of mESCs as previously described^54,55^. Briefly, we seeded clonal mESC populations onto feeder-free laminin-coated (BioLamina, Sundyberg, Sweden) tissue culture plates and maintained the cells in culture medium until cells reached confluency. We then added 200 ng/mL colchicine (Sigma-Aldrich, MO, USA) to the culture medium and incubated cells at 37°C for 3 hours. Following the incubation, we detached cells from the plate with 1:2 diluted Accumax (Thermo Fisher Scientific, MA, USA) and prepared a single-cell suspension by pipetting. We then pelleted the cells via centrifugation and subsequently resuspended the cells in 75 mM KCl. Following an incubation at 37°C for 8 minutes, we fixed cell suspensions in a 3:1 (v/v) solution of methanol and glacial acetic acid (Sigma-Aldrich, MO, USA) at room temperature. Fixed cells were dropped onto glass slides and stained with KaryoMAX Giemsa stain (ThermoFisher, MA, USA) in Gurr buffer (Gibco, MA, USA). We imaged stained cells using bright-field microscopy (EVOS M7000) and counted chromosomes using the particle analysis tool in the ImageJ software (v1.54f).

### Bionano analysis

We performed bionano analysis of mESCs according to manufacturer specifications. Briefly, we pelleted 500K to 1M mESCs and isolated ultra high molecular weight (UHMW) genomic DNA as described in the Bionano Prep SP-G2 Fresh Cell Pellet DNA Isolation Protocol (Bionano, Doc no. CG003). We then fluorescently labeled UHMW gDNA by the Direct Label Enzyme (DLE-1) and then stained for backbone visualization (Bionano Prep Direct Label and Stain (DLS) Protocol, Doc no. 30206). We loaded the labeled and stained UHMW gDNA sample onto a Saphyr Chip and into the Saphyr System Instrument (Bionano Saphyr System User Guide, Doc no. 30247). We used Bionano Solve to identify structural variations (SV) and low variant allele fraction variants (VAF) (Bionano Solve Theory of Operation: Structural Variant Calling, Doc no. CG-30110). We analyzed unedited mESCs in parallel as the background to extract SV information for edited cell lines. The resolution for detecting variants with this method is ∼500bp.

### Chimera production

We maintained mESCs in the ground state of pluripotency for up to three passages in N2B27 basal medium, modified from a previous publication^56^, seven days before 8-cell stage injection. The medium comprised of a 1:1 mixture of DMEM/F12 (Gibco, MA, USA) and Neurobasal (Gibco, MA, USA), supplemented with 1% (v/v) B-27 (Gibco, MA, USA), 0.5% (v/v) N-2 (Gibco, MA, USA), 1% (v/v) GlutaMAX (Gibco, MA, USA), 2 mM MEM Non-Essential Amino Acids (Gibco, MA, USA), 100 U/mL Penicillin-Streptomycin (Sigma-Aldrich, MO, USA), and 0.1 mM 2-mercaptoethanol (Sigma-Aldrich, MO, USA). Following Ying et al. 2008^57^, we further supplemented the medium with 1000 U/mL human LIF (Sigma-Aldrich, MO, USA) and the inhibitors 1 μM CHIR99021 (Selleckchem, TX, USA) and 0.4 μM PD0325901 (Selleckchem, TX, USA).

We produced chimeras by adapting the previously described laser-assisted protocol^16^. We cultured zygotes from C57BL/6N mice in KSOM medium (Millipore, MA, USA) for approximately 48 hours at 37°C under 5% CO₂ and 5% O₂. We selected embryos at the 8-cell stage, before compaction, for manipulation. Using a holding pipette with 10-15 μm ID (Sunlight Medical, FL, USA) to stabilize the embryos, we made a perforation in the zona pellucida (ZP) with an 800 ms tangential laser pulse (100% power) from an XYClone laser system (Hamilton Thorne Bioscience, MA, USA), targeting the outer margin of the ZP in a region away from blastomeres. We injected eight mESCs into the perivitelline space using an injection pipette with 15 μm ID (Sunlight Medical, FL, USA). Following injection, we washed the embryos and cultured them in KSOM under the same environmental conditions. Embryos that developed into blastocysts were collected and transferred to the uterine horns of pseudopregnant mice.

### Guide RNA and HDR donor design

For Cas9 editing, we designed most synthetic single-guide RNA (sgRNA) using the CRISPR Predict tool in Geneious Prime (Geneious, 2024.0.7) and others we manually designed. We assessed guide sites for CRISPR activity using established efficiency scores^58^. We evaluated off-target effects against selected regions of interest and the entire *Mus musculus* genome, GRCm39, with a tolerance of down to 3 mismatches. We further evaluated potential off-target edits in the Integrated DNA Technologies (IDT) CRISPR-Cas9 guide RNA design checker (IDT, IA, USA). We designed ssODN templates for HDR symmetrically around the cut site disrupted by the sgRNA. We used homology arms of thirty, forty, and sixty base pairs and PAM silencing mutations for some donors.

For cytosine base editing (CBE), we designed sgRNAs to target regions where the C-to-T conversion would either introduce a premature stop codon^41^ or disrupt a splice site^43^, both resulting in gene disruption and loss of function. The editing designs in this study ensured that any C-to-T bystander effect from CBE would not impede stop codon introduction or splice site mutation. However, edits other than C-to-T^59^, although less frequent, could abolish the effect of the intended point mutation. Unintended edits were considered and evaluated during guide design and genotyping.

The sequences of all sgRNA and HDR donors used in this study are listed in **Tables S4 and S5**.

### Plasmids

All plasmids were assembled via NEBuilder HiFi DNA Assembly (New England Biolabs (NEB), MA, USA). Correct assembly was verified using Oxford Nanopore whole plasmid sequencing (Plasmidsaurus, OR, USA). For zygote editing, we cloned an *in vitro* transcription plasmid template for CBE mRNA production. This vector comprised a T7 promoter to initiate transcription, CBE-P2A-mScarlet3, and flanking UTR sequences. For mESC editing, we cloned a CBE-P2A-mScarlet3 plasmid for base editor expression, four-guide array plasmids for guide RNA driven by independent U6 promoters, a BEAR reporter plasmid for base editing enrichment as previously described^48^, and a eGFP integration piggyBAC plasmid for generating a testing cell line of the high yield chimera workflow. We used Tad-CBEd^59^ with a spRY PAM^44^ preference as the CBE of choice.

### *In vitro* transcription

To produce the CBE mRNA used for zygote editing, we generated a linear DNA amplicon that included the T7 promoter, coding regions, and UTRs using KOD One (Sigma-Aldrich, MO, USA) and a plasmid template. We performed gel electrophoresis and excised the band corresponding to the amplicon, which we gel purified using the QIAquick Gel Extraction Kit (Qiagen, MD, USA). We then bead purified the eluted DNA using AMPure XP Beads (Beckman Coulter, CA, USA).

The purified amplicon served as a template for an *in vitro* transcription reaction using the HiScribe® T7 High Yield RNA Synthesis Kit (NEB, MA, USA). We incubated the reaction at 37°C for 2 hours, and then purified mRNA using the Monarch RNA Cleanup Kit (NEB, MA, USA). We measured the yield and purity of the mRNA using a Denovix DS-11 (Denovix Inc, DE, USA) and aliquoted purified mRNA, which we stored at -80°C until use.

### Genotyping

To collect samples for genotyping, we harvested embryos at the blastocyst stage, collected mESCs from single-cell Accumax (Thermo Fisher Scientific, MA, CA) suspensions, and obtained ear notch samples from individual mice 21–26 days after birth. We lysed each sample with Quick Extract (Fisher Scientific Company, NH, USA) and placed each in a thermocycler at 65°C for 15 minutes followed by 98°C for 2 minutes.

We amplified genomic regions spanning the mutation of interest via PCR, followed by next generation sequencing (NGS) or Sanger sequencing as described below. All primer sequences used for genotyping are listed in **Table S6** and were designed in Geneious Prime (Geneious, 2024.0.7).

To prepare amplicons for NGS genotyping, we performed two step PCR. PCR1 used KOD ONE (Sigma-Aldrich, MO, USA) and a thermocycle consisting of 30-35 cycles of 98°C for 15 seconds, 60°C for 5 seconds, and 68°C for 5 seconds. We added 1 µl of PCR1 into the PCR2 mixture that included custom indexes for i5 and i7 adapter sequences (IDT, IA, USA). The PCR2 thermocycle consisted of 8 cycles of 98°C for 15 seconds, 60°C for 5 seconds, and 68°C for 5 seconds. We confirmed amplification via gel electrophoresis with 5 µl of PCR2.

Next, we pooled PCR2 samples into a purified library pool using AMPure XP Beads (Beckman Coulter, CA, USA) at 1:1 ratio, quantified DNA concentration by both Qubit Flex Fluorometer (Thermo Fisher Scientific, MA, USA), and visualized library amplicon size with a D1000 ScreenTape (Agilent Technologies, CA, USA) on the Agilent TapeStation. We sequenced libraries using either a 300- or 500-cycle kit on the Illumina MiSeq or the NextSeq2000 instruments (Illumina, CA, USA) targeting ten thousand reads per sample. We uploaded the data onto the Form Bio (Form Bio Inc., TX, USA) platform and ran the VALID workflow. Briefly, sequence read pairs are merged using fastp^60^, then reads are aligned to the expected amplicons using Magic-blast^61^ for edited and wild-type cells. We used these alignments to calculate editing efficiencies and identify off-target edits overlapping the amplicon region.

For Sanger sequencing, we performed 15 µl PCRs using KOD ONE (Sigma-Aldrich, MO, USA) as follows: 38-40 cycles of 98°C for 15 seconds, 60°C for 5 seconds, and 68°C for 5 seconds. After PCR was complete, we ran 5 µl of the samples on a 2% agarose gel to confirm presence of a band. We sent the remaining 10 µl of the unpurified PCR to Genewiz/Azenta (Genewiz/Azenta, NJ, USA) for Sanger sequencing. We ran ab1 files from Sanger sequencing on the Synthego website for ICE Analysis.

We define gene knock-out (KO) as exonic insertions or deletions (indels) not in multiples of three, an exonic indel over 100 bp, introduction of a stop codon, or a splice site mutation. For large indels that may influence the mapping of NGS reads (usually >100 bp), we used electrophoresis band patterning and Sanger sequencing to confirm absence of wild-type sequences. All mice were genotyped primarily by NGS and supplemented by band patterning and Sanger sequencing.

### Whole genome sequencing and off-target analysis

For each sample, we extracted genomic DNA using the NucleoSpin Tissue kit following manufacturer instructions (Macherey-Nagel, Düren, Germany). Libraries were prepared using the Illumina DNA Prep kit (Illumina, CA, USA). We used double-sided bead purification to purify amplified libraries which we quantified using an Agilent TapeStation (Agilent Technologies, CA, USA) prior to normalization and pooling. Libraries were then sequenced on the NextSeq2000 (Illumina, CA, USA) using XLEAP-SBS chemistry targeting 30x genomic coverage per sample.

We identified editing at off-target sites by whole genome sequencing and alignment. We used the Valid WGS workflow on the Form Bio platform to identify SNVs, indels, and structural variants to identify edits. In this workflow, we trimmed reads using fastp and then aligned them to the mm39 reference genome^62^ using Sentieon optimized BWA MEM^63,64^. Duplicate reads were marked using Picard MarkDuplicates. Somatic variants were detected using the Sentieon TNHaplotyper2 tool while structural variant calls were detected using DNAscope. We filtered variants based on multiple filtering criteria (Strand bias, Alternate Allele Count < 2, Variant Allele Frequency < 0.7) to predict on and off target edits. After filtering, on target and off target variants list were split based on the intended location of on targets. Quality reports were produced by MultiQC. Variant effects were determined using snpEff.

For zygote editing, we took a targeted approach to determine off-target editing to account for normal genetic variations between individuals. We predicted off-target binding sites for all multiplexed guides in each zygote-editing experiment using the Form Bio platform (Formbio, Texas, USA) via the DR GENE workflow, which uses an in-house implementation of the runCrisprBwa function from the crisprVerse package in R^65^. All predicted off-target sites with three or fewer mismatches to the sgRNA protospacers were interrogated for off-target editing from the WGS data. For mice produced from CBE-edited zygotes, variants found within ±20bp of a predicted off-target binding site were considered off-target edits. To account for larger potential alterations due to NHEJ in the mice produced from Cas9-edited zygotes, variants found within ±200bp of a predicted off-target binding site were considered off-target edits. Of these, all variants with a VAF greater than 0.7 and read depth greater than 5 are reported. As ESC and ESC derived samples originate from the same B6129 ESC cell line, we report all off-target variants with a variant allele frequency (VAF) greater than 0.7 and read depth greater than 5.

## Supplemental Figures

**Figure S1.**
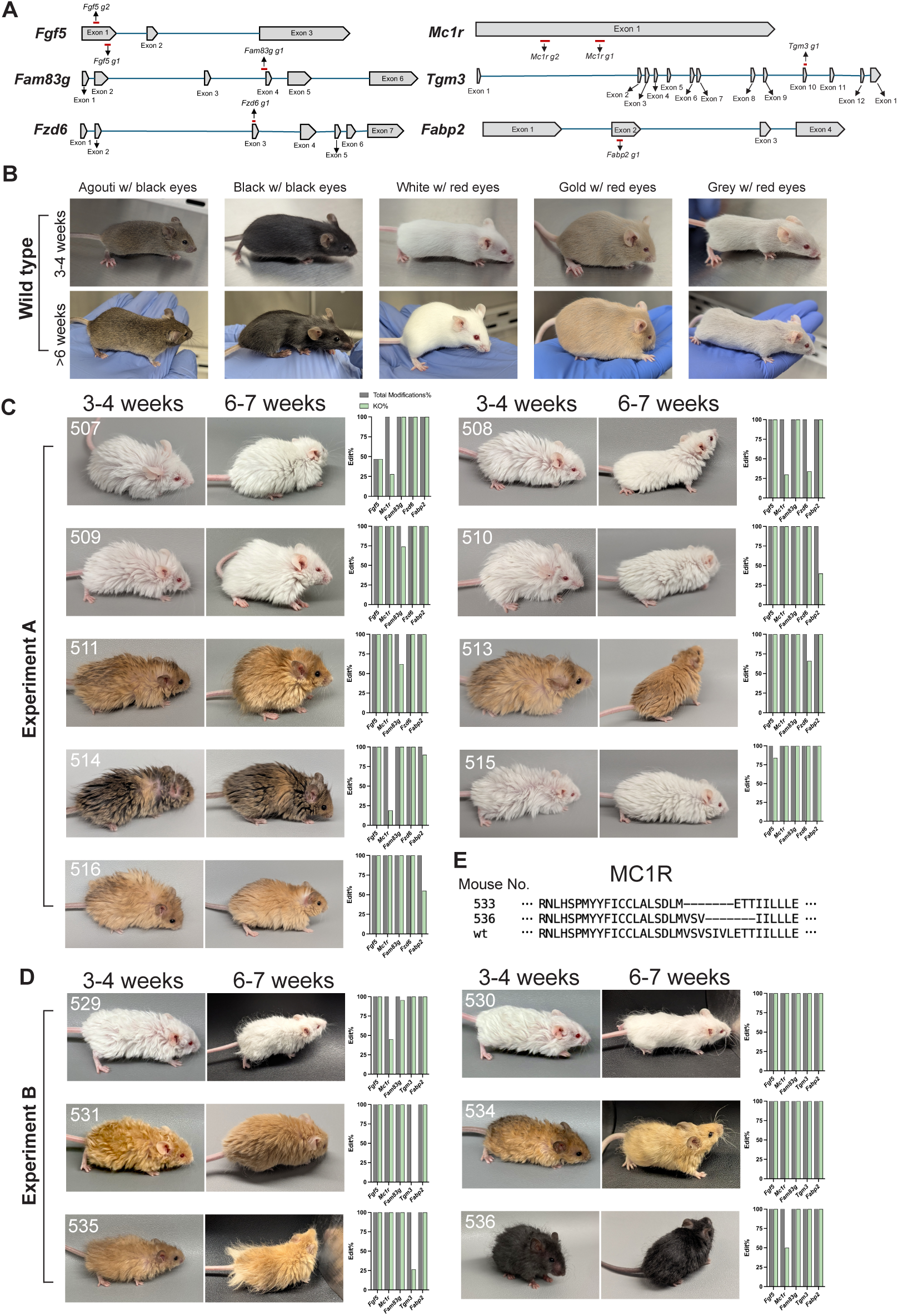
Multiplex Editing of Zygotes with CRISPR/Cas9, related to. Figure 2**. (A)** Target genes and the location of their sgRNAs chosen for Cas9 KO animal production. We selected production sgRNA according to editing efficiency, off-target prediction, and the location of the edit. The ideal edit location should either mimic the previous literature or be in an early exon to facilitate KO. **(B)** Wild-type B6SJL F2 mice (B6 × SJL)F1 × (B6 × SJL)F1 have a variety of coat+eye color combinations: agouti with black eyes (the most common), black with black eyes, white with red eyes, gold with red eyes (rare), and diluted grey with red eyes (rare). Two age points are shown here. The agouti mouse pictures are the same as used in Fig. 2 and Fig. 3. **(C, D)** The editing profiles and photos of mice not shown in Fig. 2 from Cas9 KO Experiment A and B. **(E**) Alignment of the two different types of 7-amino-acid deletions in MC1R (from mouse #533 and #536) with the wild-type MC1R protein sequence.

**Figure S2.**
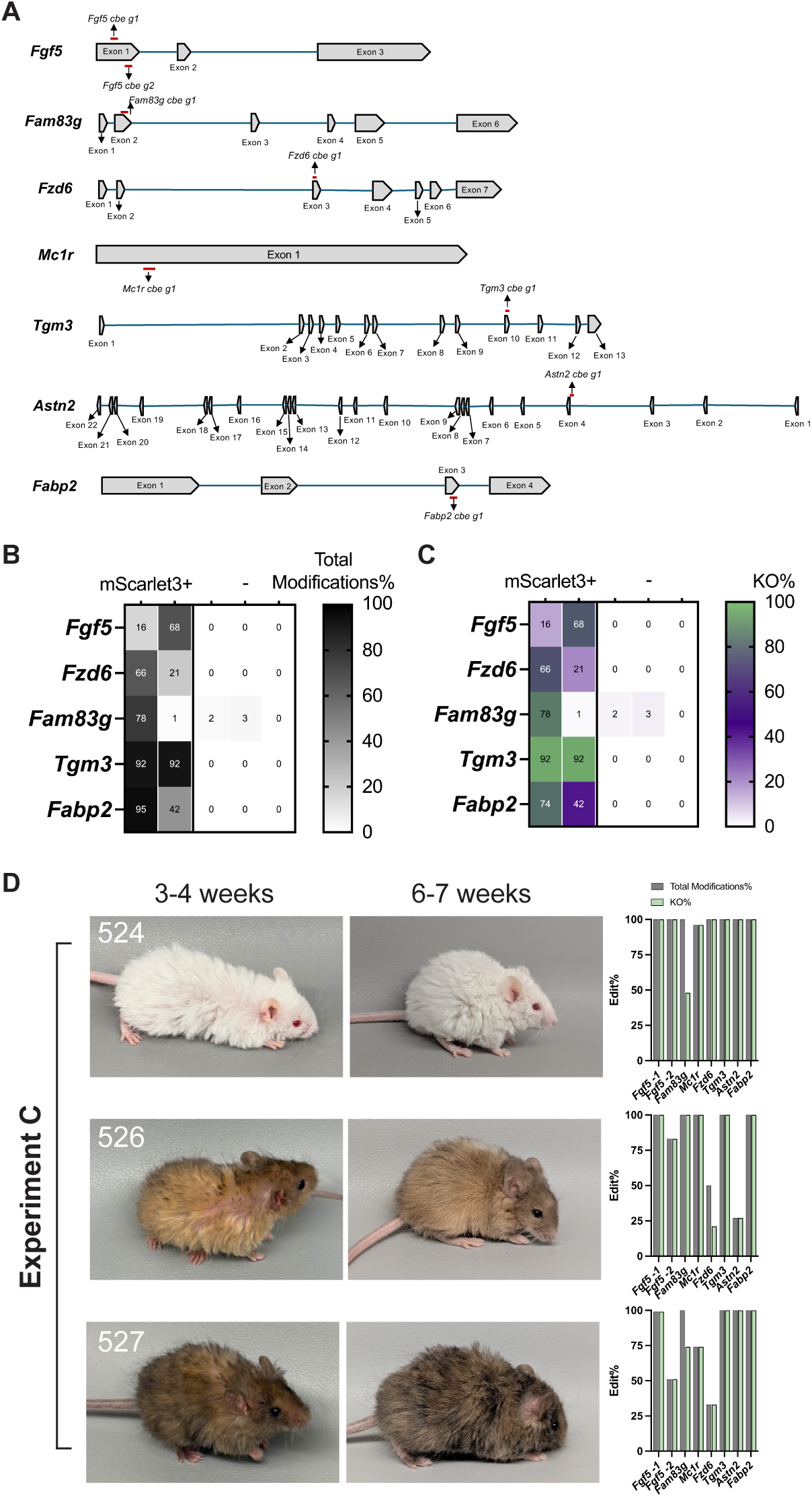
Multiplex Editing of Zygotes with CBE, related to. Figure 3**. (A)** Target genes and the location of their sgRNAs chosen for CBE KO animal production. We selected animal production sgRNA according to editing efficiency, off-target prediction, and the location of the C-to-T mutation. The ideal edit location should either mimic the previous literature or be in an early exon to facilitate KO. **(B, C)** Single embryo total modification% and KO% comparison between mScarlet3 + and mScarlet3 - embryos demonstrated in Figure 3C. The individualized data represent the multiplexability and editing mosaicism in one animal by each condition. Each column is data from 1 embryo. Editing was determined by Sanger sequencing. **(D)** The editing profiles and photos of mice not shown in Figure 3 (Experiment C).

**Figure S3.**
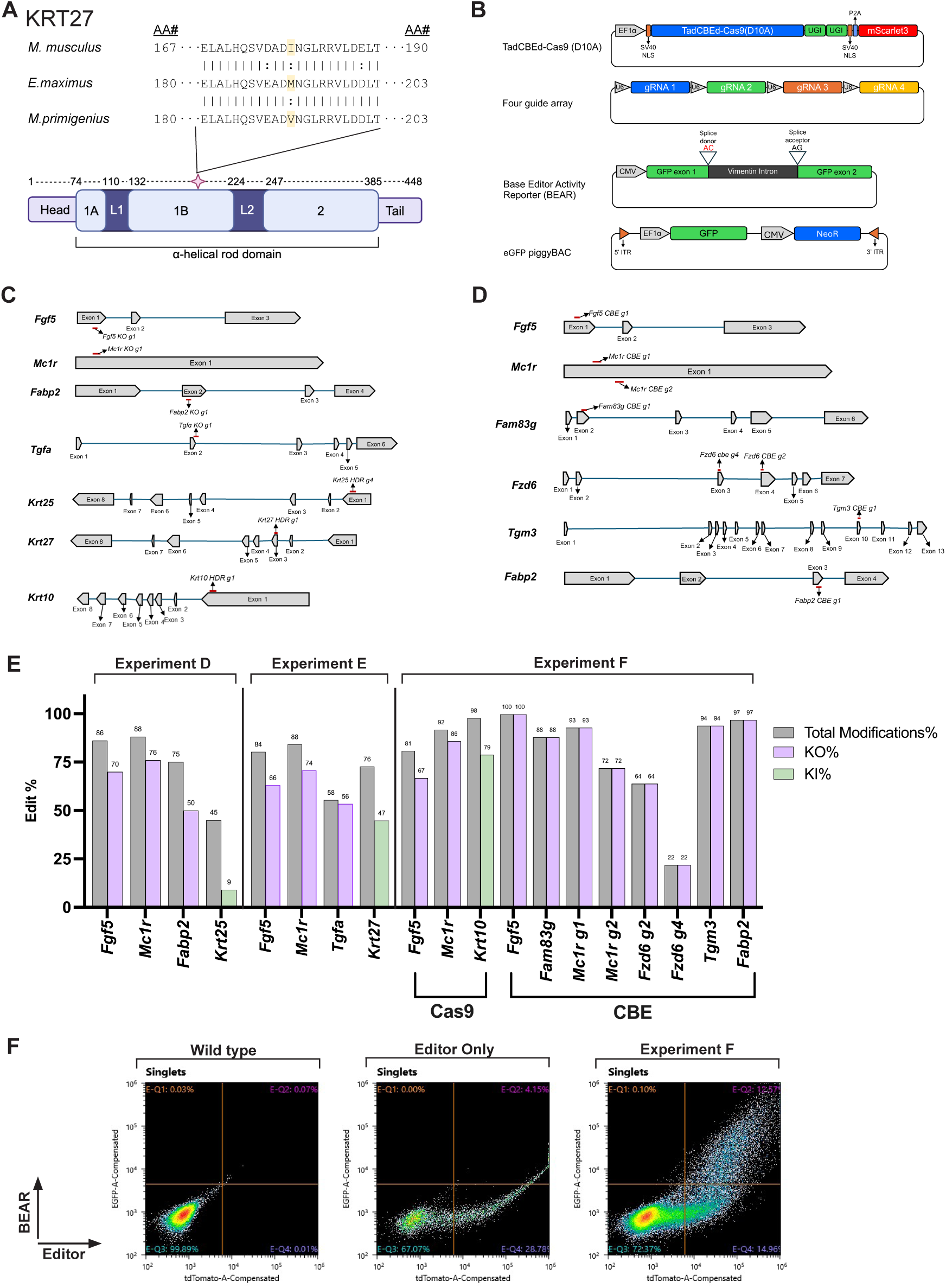
Multi-modality multiplex editing in mESC, related to. Figure 4**. (A)** KRT27 protein alignment of *Mus musculus, Elephas maximus,* and *Mammuthus primigenius* centered around amino acid (AA) 191 (highlighted in yellow), a fixed coding mutation between *E. maximus* (methionine) and *M. primigenius* (valine). In Experiment E, the *M. musculus* isoleucine was edited to *M. primigenius* valine via HDR. This mutation occurs in the 1B ɑ-helical rod domain of the secondary structure of the KRT27 protein (star). **(B)** Maps of plasmids used in Experiment D, E, and F. **(C)** Target genes and the location of the sgRNAs selected for Cas9 KO and Cas9 HDR mESC generation. **(D)** Target genes and the location of the sgRNAs selected for CBE KO mESC generation. **(E)** The KO% and KI% from Experiment D, E, and F bulk mESC populations prior to single-cell sorting. **(F)** Gating strategy for single cell sorting Experiment F mESCs. First, we applied a size gate to select mESCs and remove iMEF carryover, followed by another size gate to exclude doublets. In the singlet gate, a quadrant gate was created to divide the diagram into four quadrants named Q1, Q2, Q3, Q4. Experiment F mESCs falling in Q2 (CBE editor (+) and BEAR (+)) were selected for single cell isolation.

**Figure S4.**
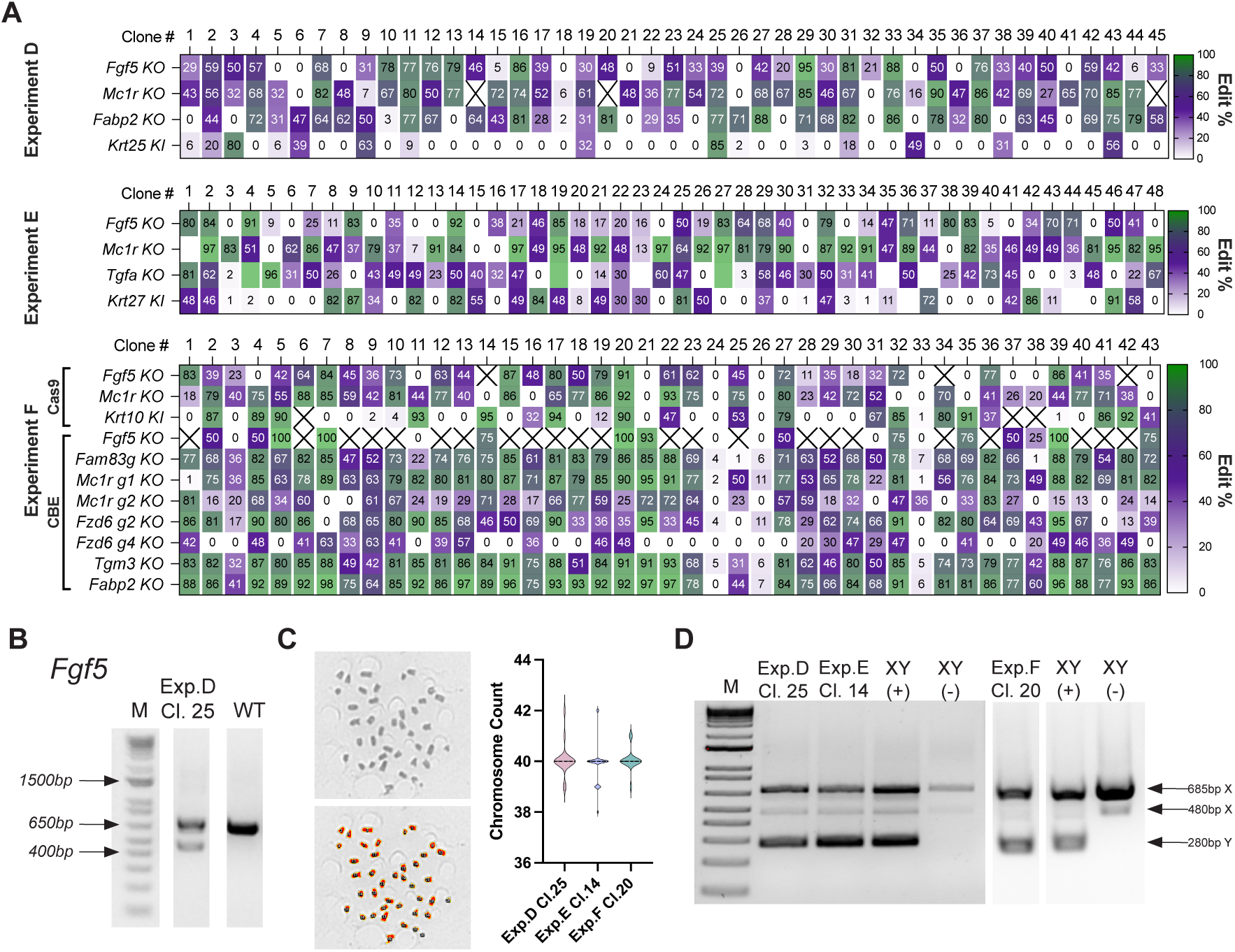
Clonal isolation of highly edited mESCs with normal chromosomal profiles, related to. Figure 4**. (A)** Preliminary KO and KI editing profiles for 45 clones generated in Experiment D, 48 clones generated in Experiment E, and 43 clones generated in Experiment F. We collected clones from an iMEF feeder layer and genotyped rapidly by Sanger sequencing. “X” marked the genotyping dropouts which was uninterpretable from Sanger results. The remaining data was sufficient to determine high value clones for further validations. **(B)** PCR analysis of *Fgf5* from Experiment D, clone 25, which shows a large deletion that was not initially identified by ICE analysis. The DNA ladder marker is labeled as “M”. **(C)** Representative mESC metaphase spread used for chromosome counting. Images were acquired using an EVOS M7000 with 20X objective. Upper left is the original photo (greyscale); lower left is the same photo where chromosomes were recognized (highlighted by colors) and counted by ImageJ. We counted chromosomes from n = 20 (Experiment D), n = 29 (Experiment E), and n = 17 (Experiment F) edited mESCs. Experiment D, clone 25, was 80% normal with an average count of 40.1 chromosomes and median count of 40 chromosomes. Experiment E, clone 14, was 70% normal with an average count of 39.90 chromosomes and median count of 40 chromosomes. Experiment F, clone 20, was 83% normal with an average count of 40.1 chromosomes and median count of 40 chromosomes. **(D)** XY PCR of Experiments D/E/F edited mESCs used for animal generation. The positive control (+) and negative control (-) are mESC lines characterized previously for internal use.

**Figure S5.**
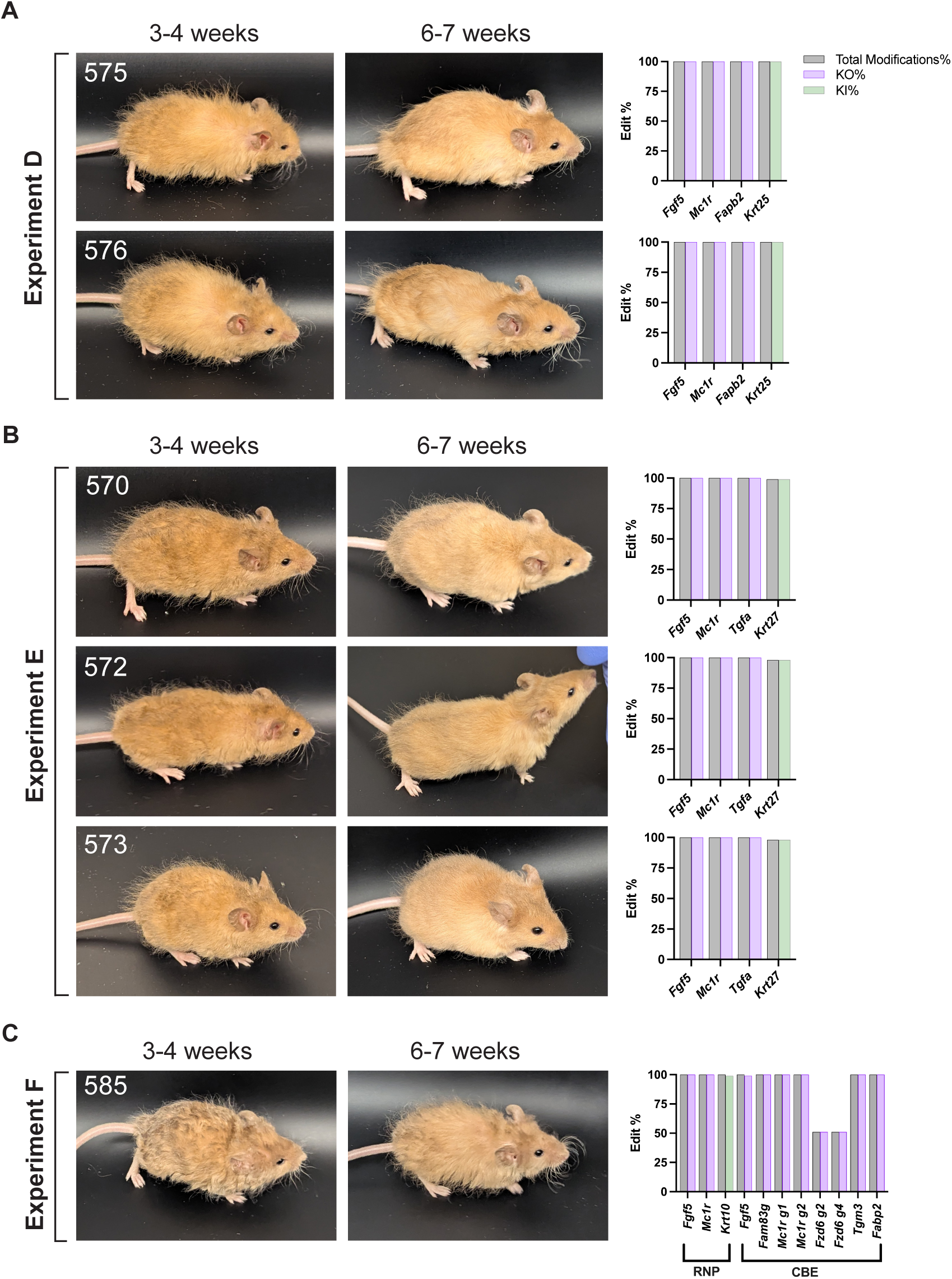
Near full mESC contribution mice produced by high yield chimera, related to. Figure 5**. (A-C)** Editing profiles and photos of mice generated from Experiment D, E, and F mESCs not shown in Figure 5.

**Figure S6.**
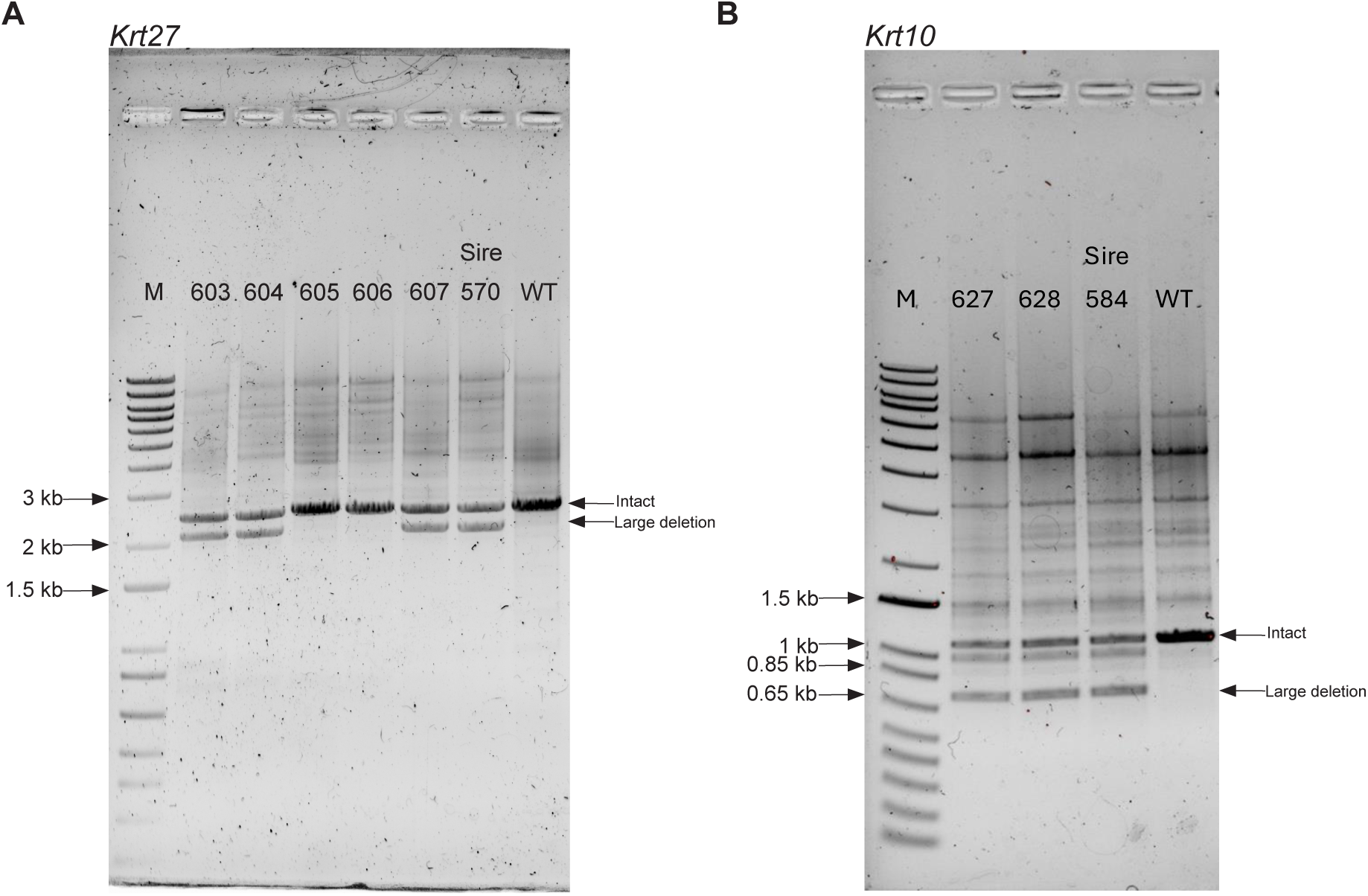
Previously undetected large deletion from Cas9 HDR editing and hemizygosity inheritance by the F1 progenies, related to. Figure 6. PCR analysis of *Krt27* and *Krt10* genotypes on the F0 sire and WT cross-breeding progeny from Experiment E and F. The sires (#570 for *Krt27*; #584 for *Krt10*) show a large deletion on one allele that was not discovered by NGS before the breeding scheme. The large deletion also showed up in some progenies (#603, #604, and #607 for *Krt27*; #627 and #628 for *Krt10*), correlating with their lack of *Krt27 KI* or *Krt10 KI* allele in Figure 6D. On the contrary, progeny #605 and #606 exhibited one normal size band, correlating with the heterozygous *Krt27 KI* genotype. The DNA ladder marker is labeled as “M”.

## Supplemental Tables

**Table S1.** Trait engineering target genes in this study, related to Figure 1.

| Gene | Expected Phenotype | Edit type | Modality used | Workflow |  | Experiment |
| --- | --- | --- | --- | --- | --- | --- |
| *Astn2* | Ridge hair | Exon-skip | CBE | Zygote editing |  | C |
| *Fam83g* | Woolly hair | KO | Cas9; CBE | Zygote editing | High efficiency chimera | A B C F |
| *Fzd6* | Disoriented hair | KO | Cas9; CBE | Zygote editing | High efficiency chimera | A C F |
| *Tgm3* | Curly hair | KO | Cas9; CBE | Zygote editing | High efficiency chimera | B C F |
| *Fgf5* | Long hair | KO | Cas9; CBE | Zygote editing | High efficiency chimera | A B C D E F |
| *Mc1r* | Gold hair | KO | Cas9; CBE | Zygote editing | High efficiency chimera | A B C D E F |
| *Fabp2* | Lipid metabolism; Mammoth-motivated | KO | Cas9; CBE | Zygote editing | High efficiency chimera | A B C D F |
| *Tgfa* | Wavy hair | KO | Cas9 |  | High efficiency chimera | E |
| *Krt25* | Curly hair | KI | Cas9 HDR |  | High efficiency chimera | D |
| *Krt27* | Textured hair; Mammoth-motivated | KI | Cas9 HDR |  | High efficiency chimera | E |
| *Krt10* | Wavy hair | KI | Cas9 HDR |  | High efficiency chimera | F |

**Table S2.**
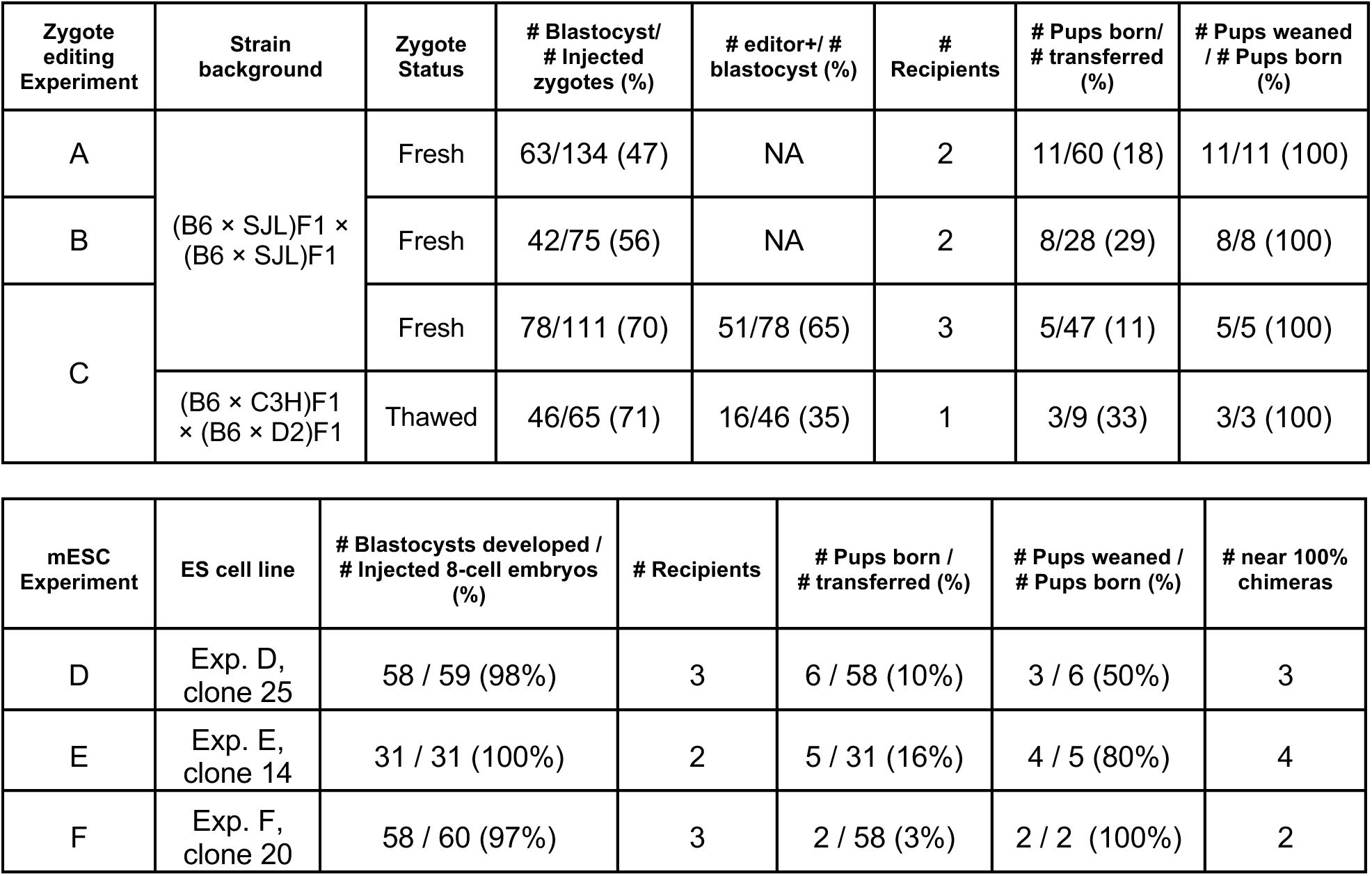
Zygote and mESC editing animal production summary.

**Table S3.** Off-target analysis summary from mESC clone unbiased WGS, related to Figure 4.

| Clone | Impact | Variant Annotation | nVariants |
| --- | --- | --- | --- |
| Exp.D-clone 25 | MODIFIER | intergenic_region | 41 |
|  | MODIFIER | intron_variant | 22 |
|  | MODIFIER | upstream_gene_variant | 8 |
|  | MODIFIER | downstream_gene_variant | 6 |
| Exp.E-clone 14 | MODERATE | missense_variant | 1 |
|  | MODIFIER | intergenic_region | 54 |
|  | MODIFIER | intron_variant | 46 |
|  | MODIFIER | upstream_gene_variant | 15 |
|  | MODIFIER | downstream_gene_variant | 17 |
|  | MODIFIER | non_coding_transcript_exon_variant | 2 |
| Exp.F-clone 20 | MODIFIER | intergenic_region | 71 |
|  | MODIFIER | intron_variant | 58 |
|  | MODIFIER | upstream_gene_variant | 23 |
|  | MODIFIER | downstream_gene_variant | 12 |

**Missense variant detail**
|  | Chr | Start | End | Gene | Ref | Alt | Annotation | VAF | Biotype |
| --- | --- | --- | --- | --- | --- | --- | --- | --- | --- |
| Exp.E-clone 14 | chr11 | 107869130 | 107869130 | *Prkca* | C | A | Missense variant | 0.72 | Protein coding |

**Table S4.**
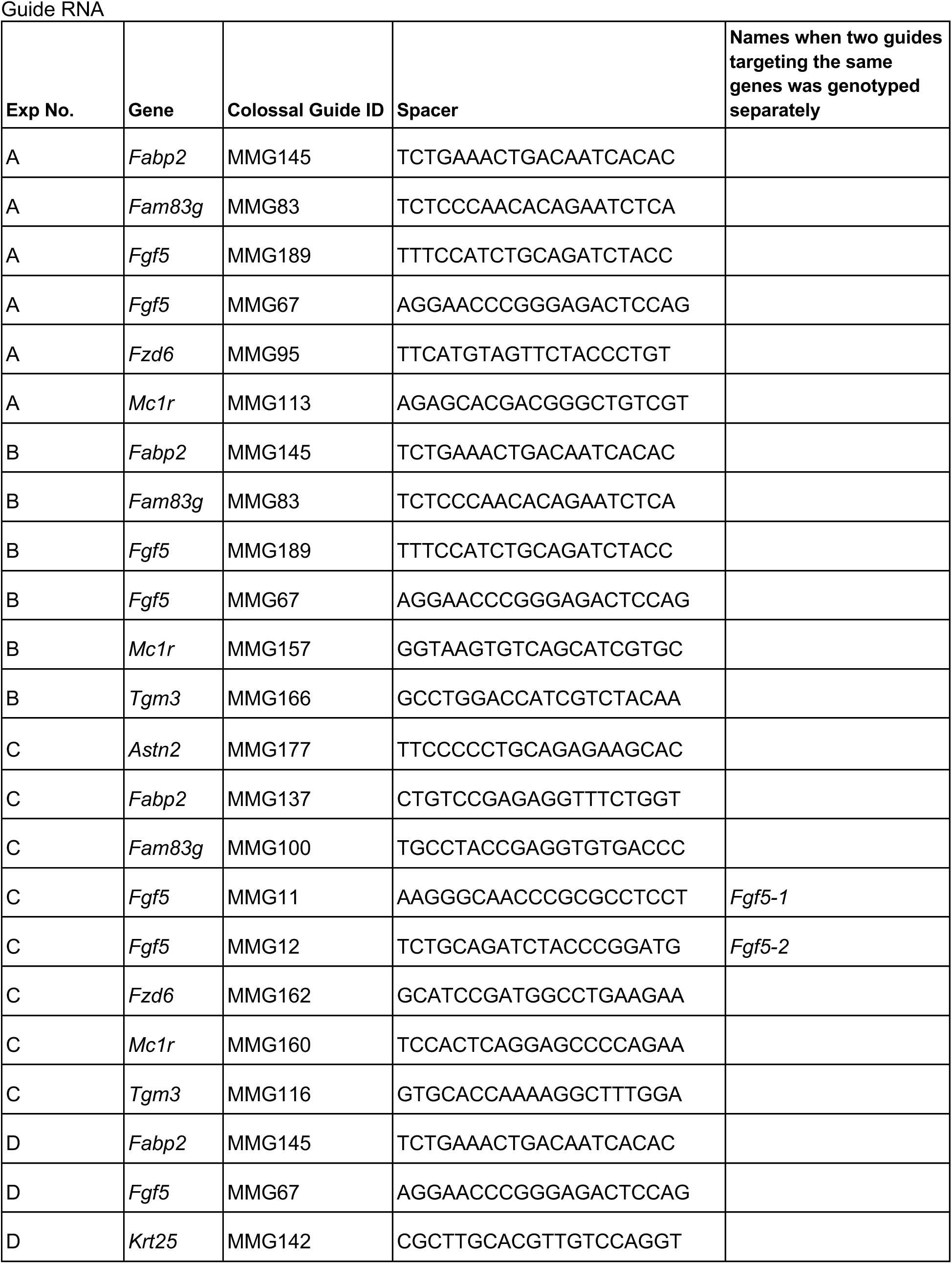
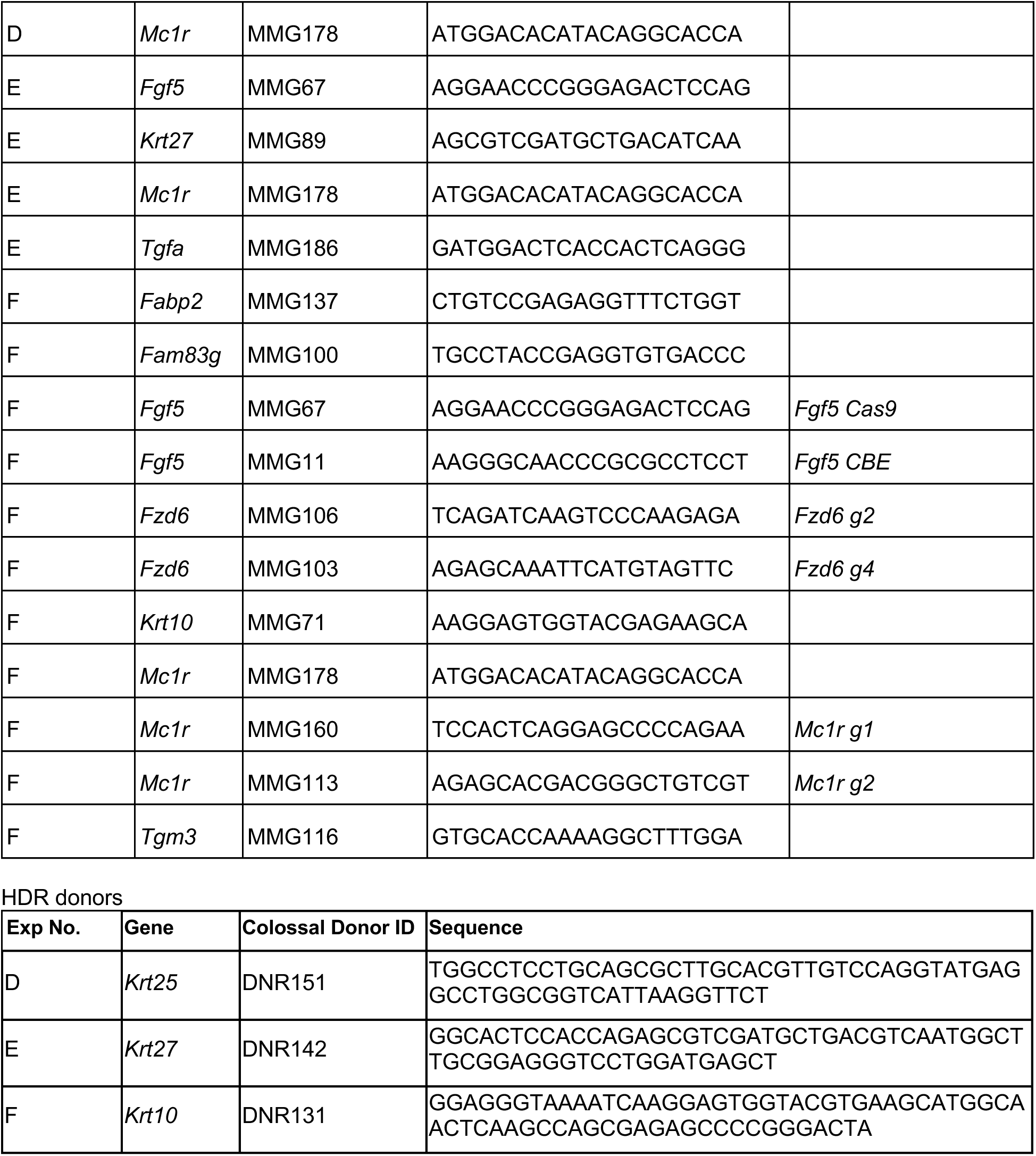
guide RNA and donor sequences used for mouse production experiments, related to Methods.

**Table S5.**
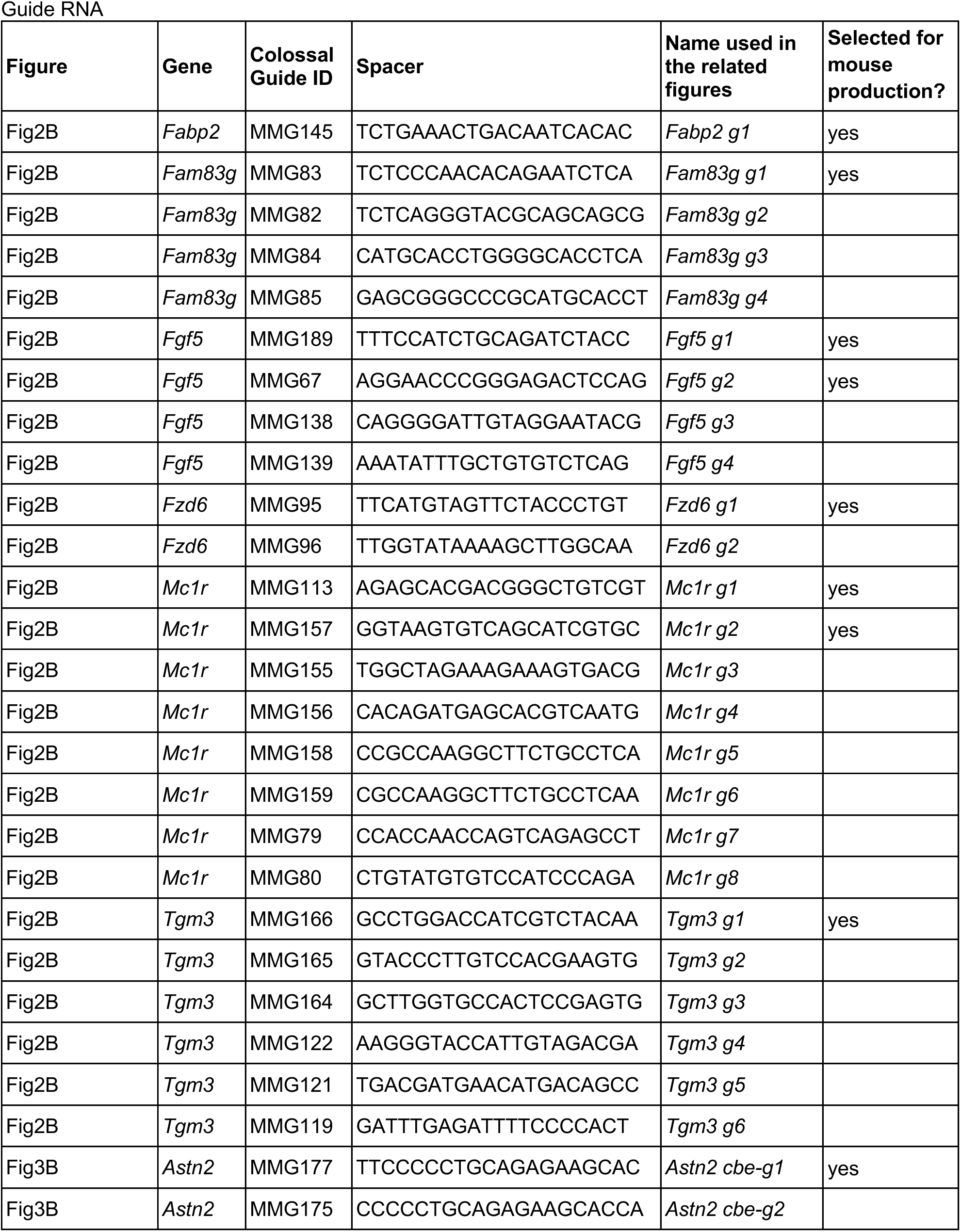
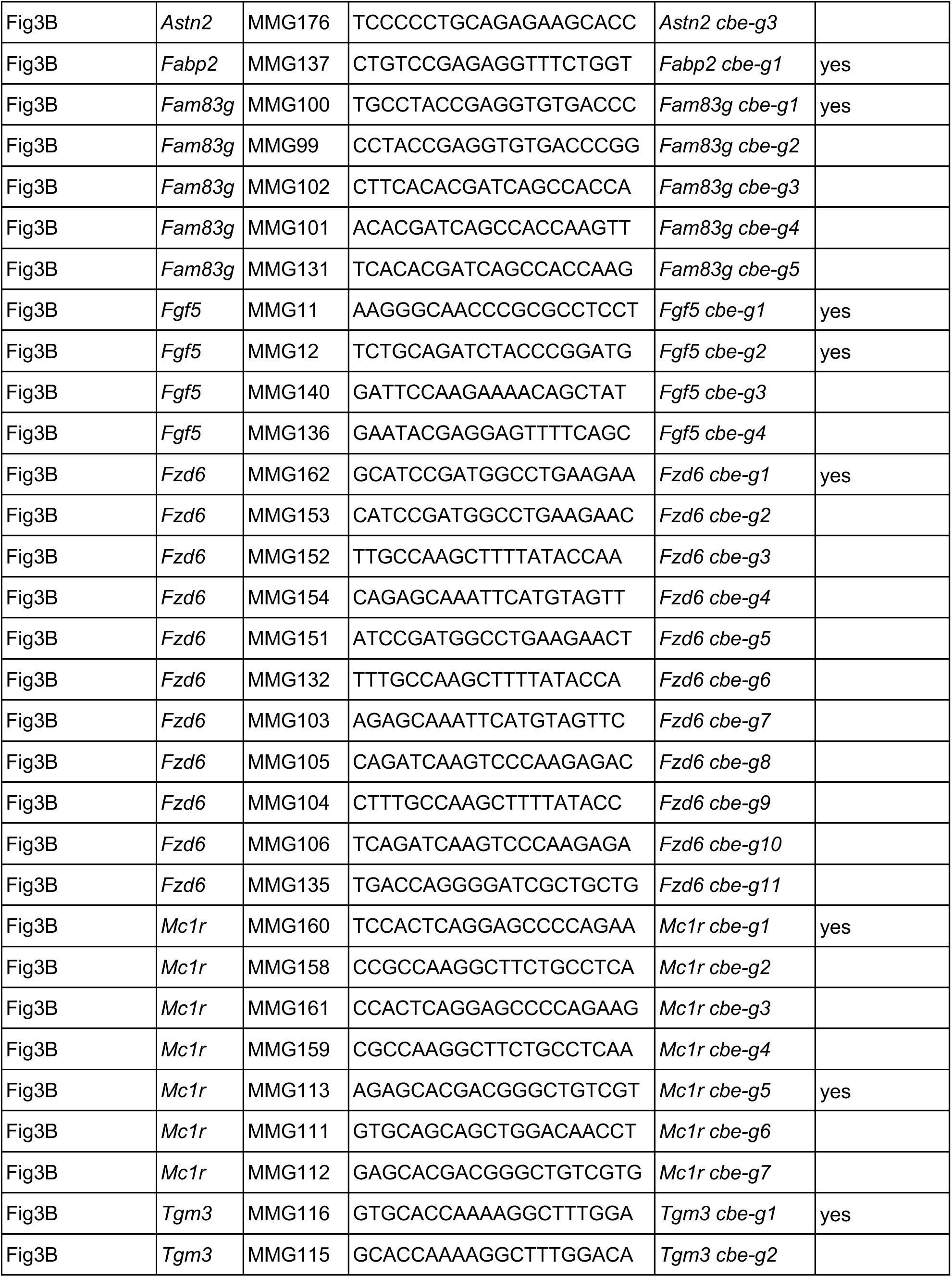
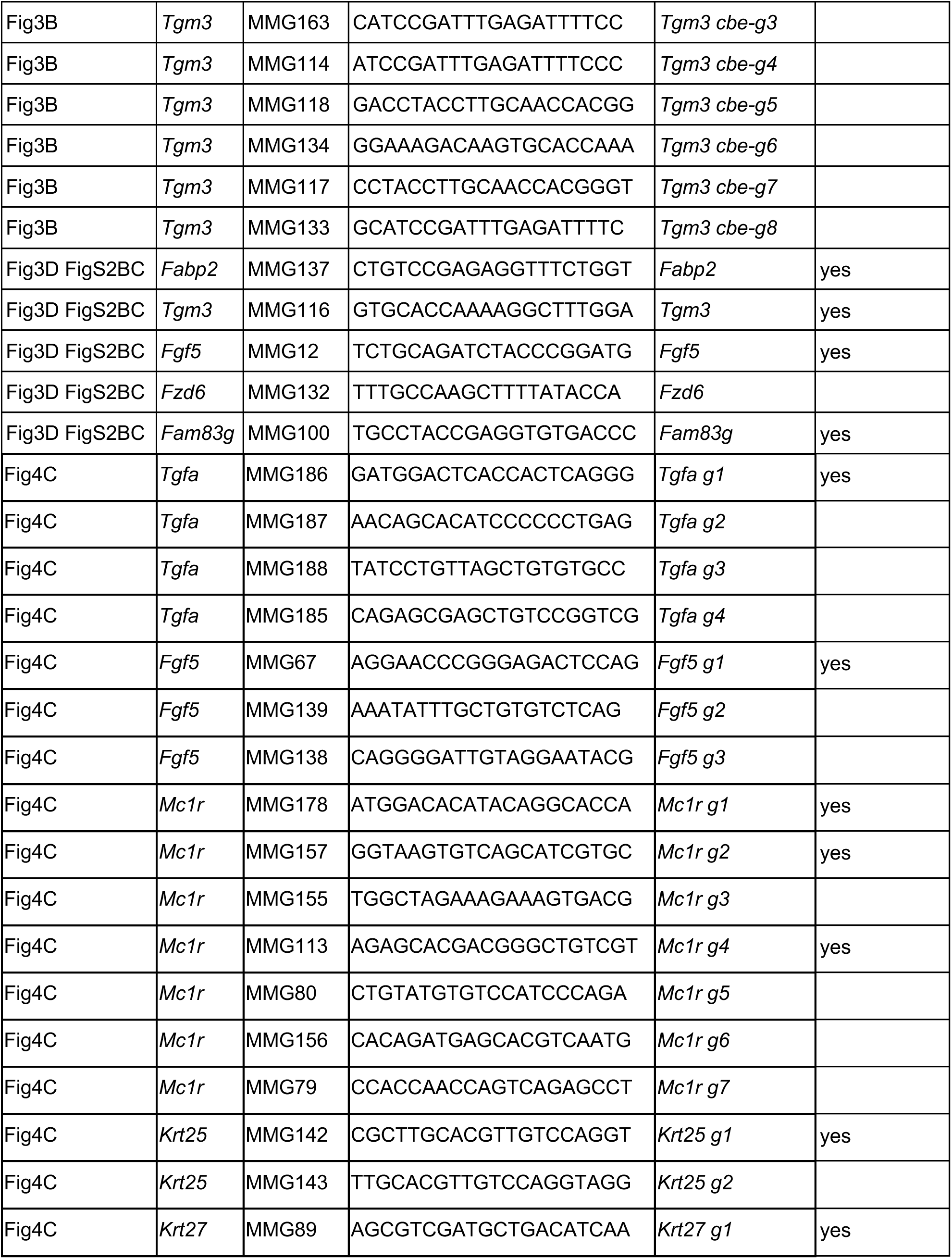
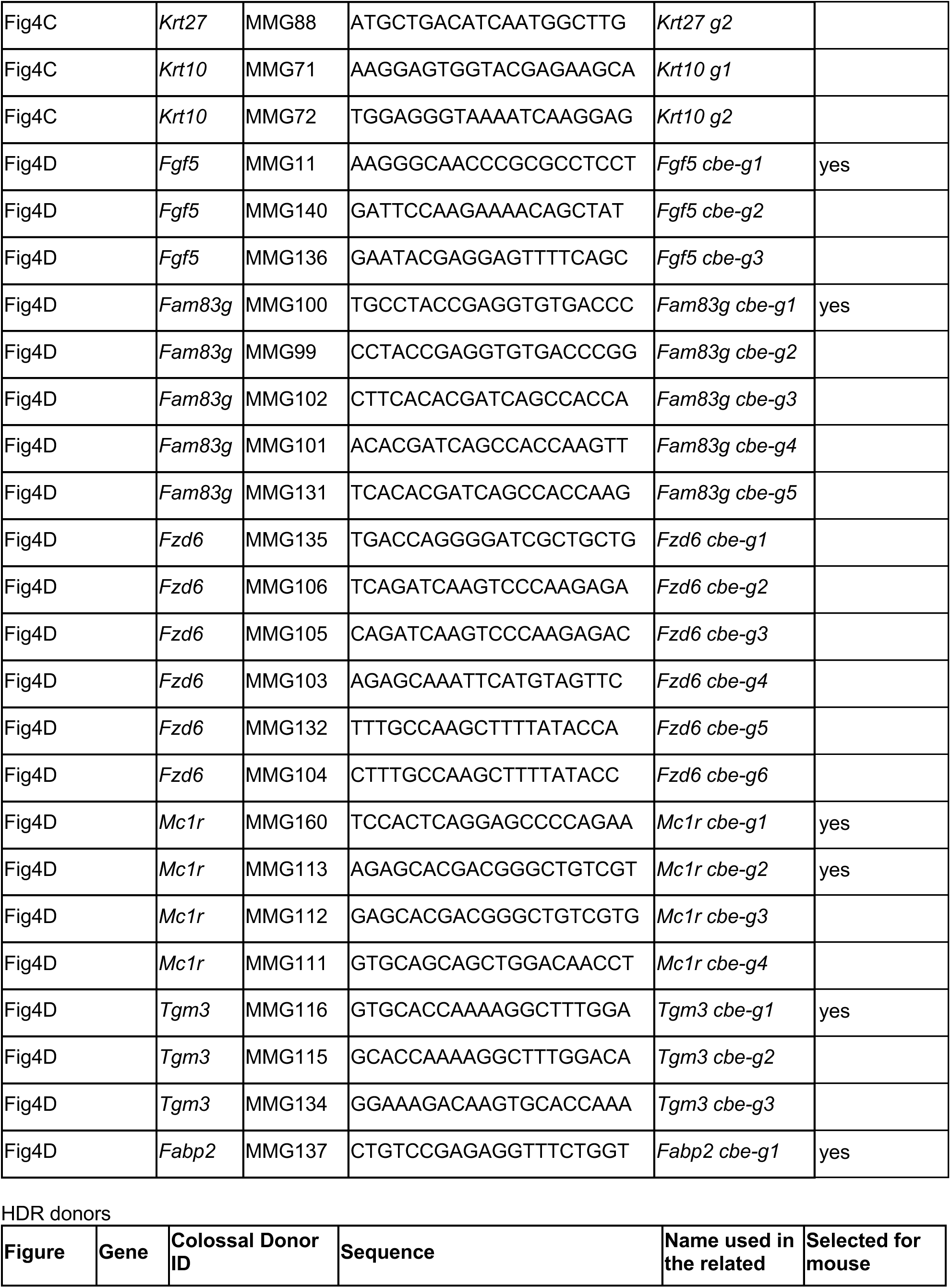
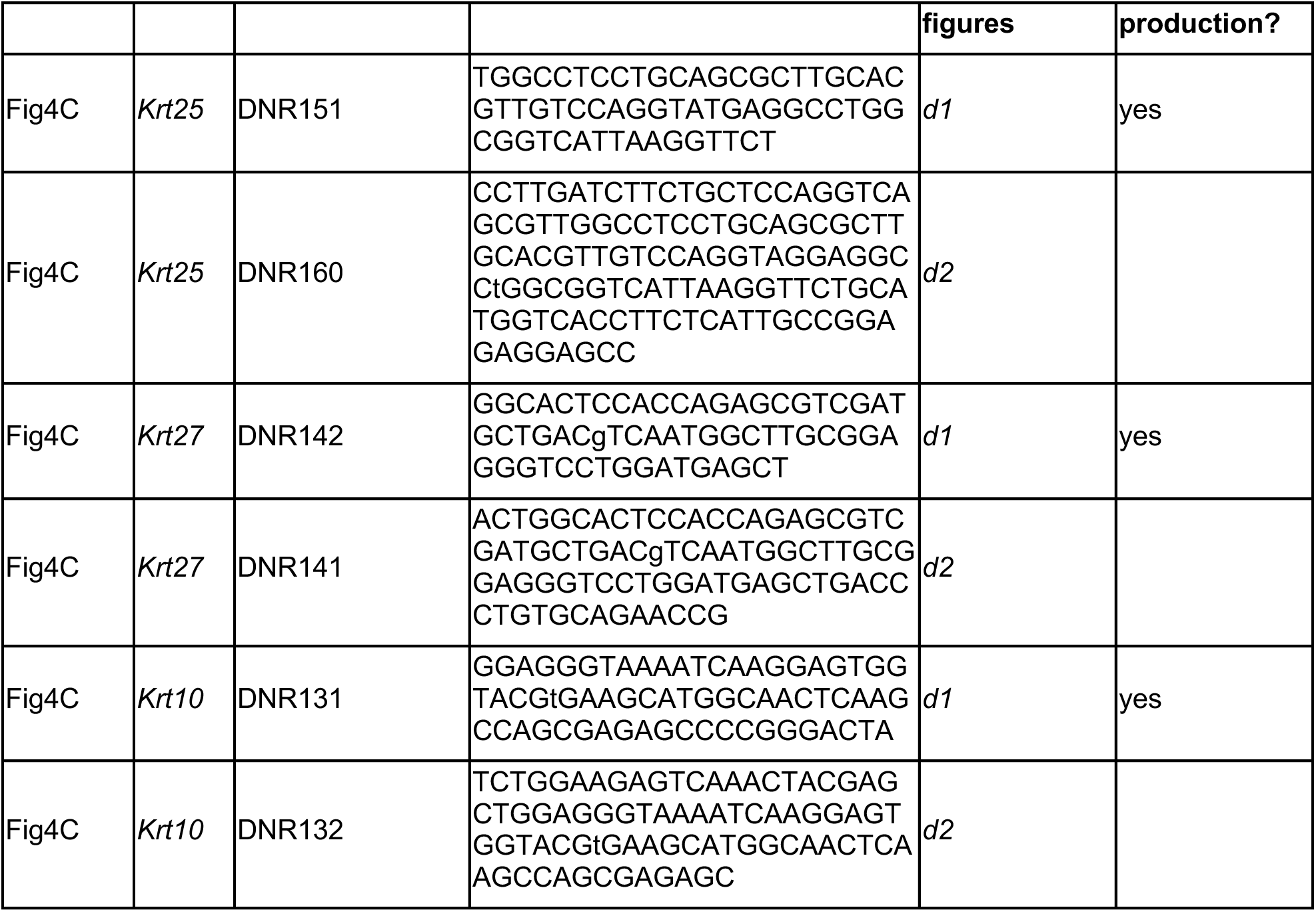
guide RNA and donor sequence used in screening and testing experiments by figures, related to methods.

**Table S6.**
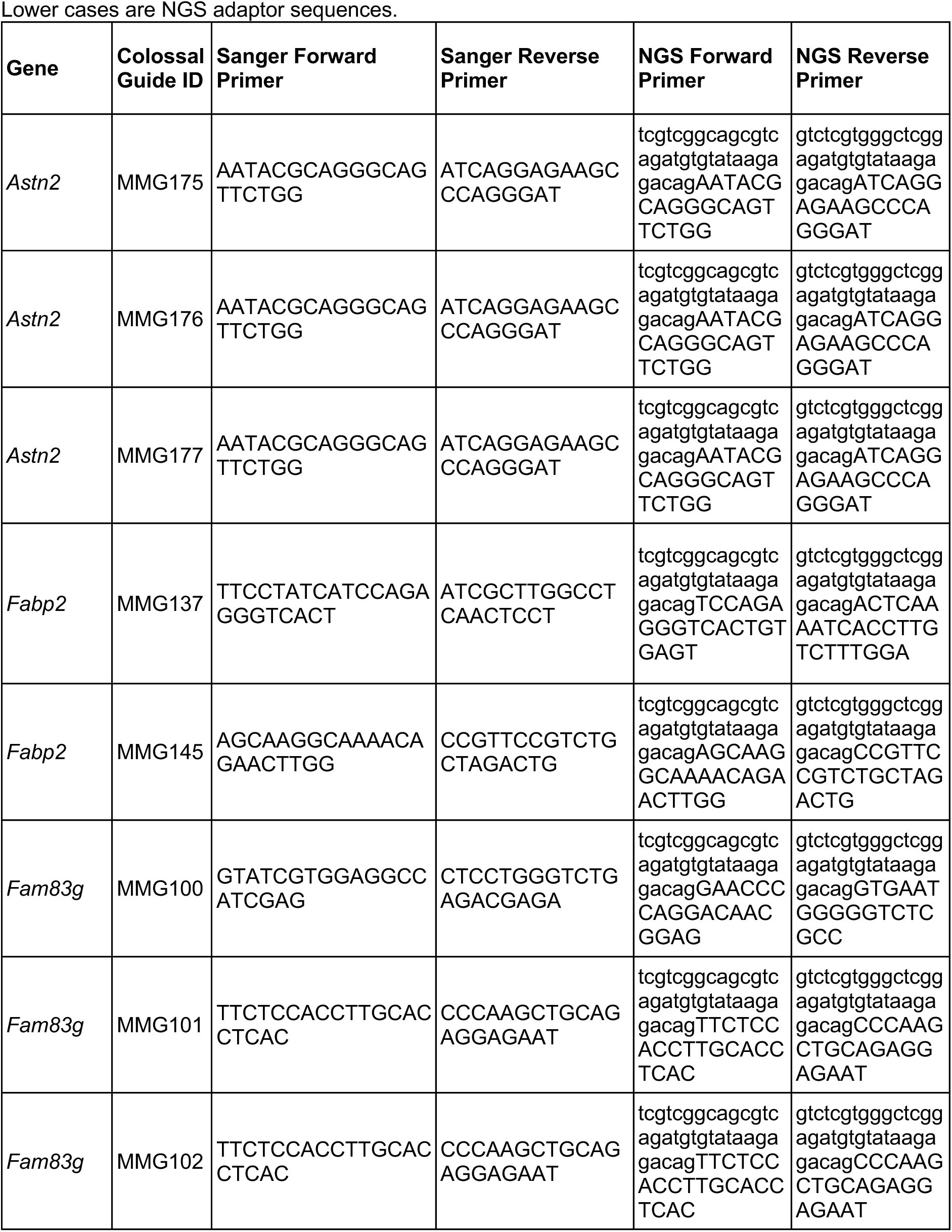
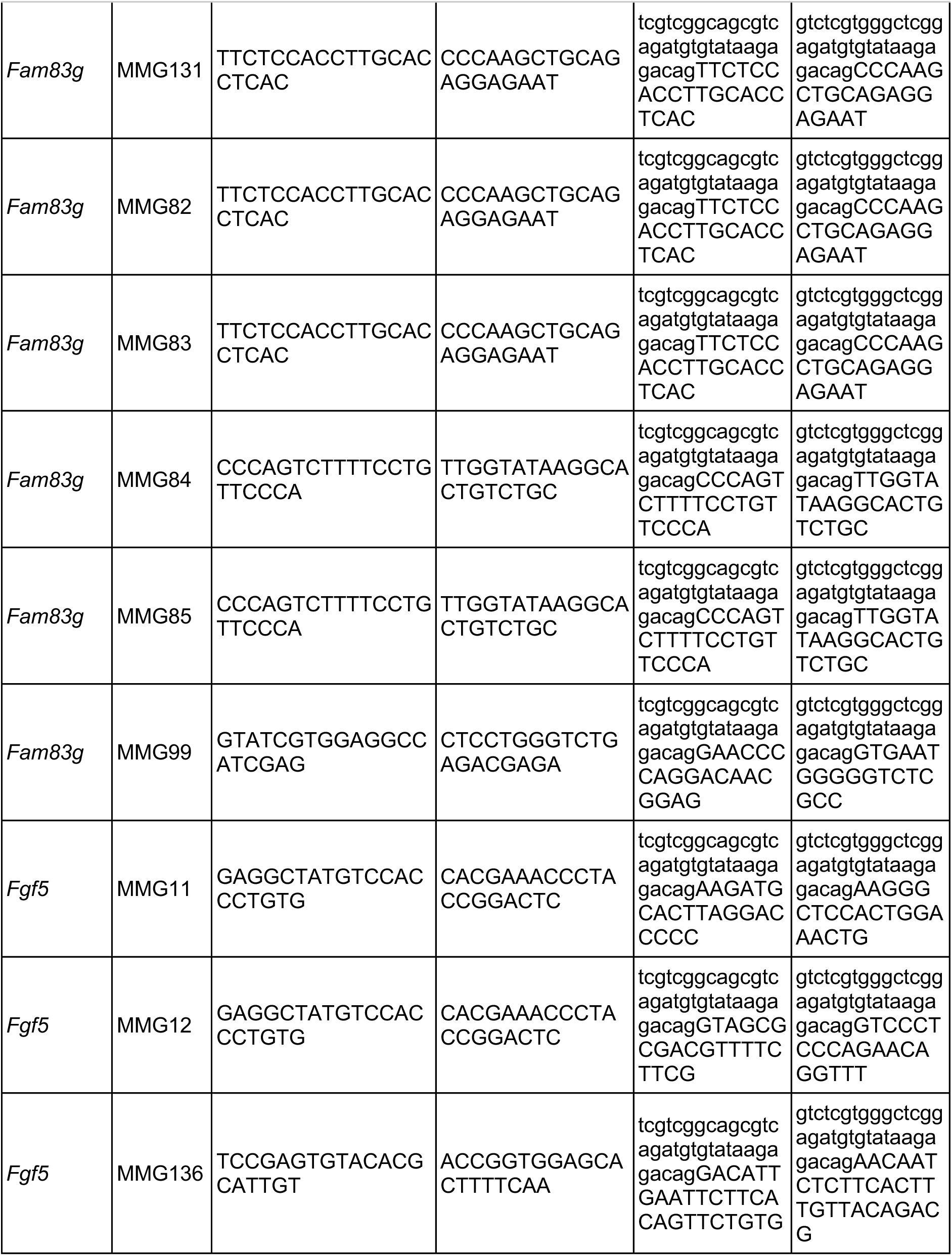
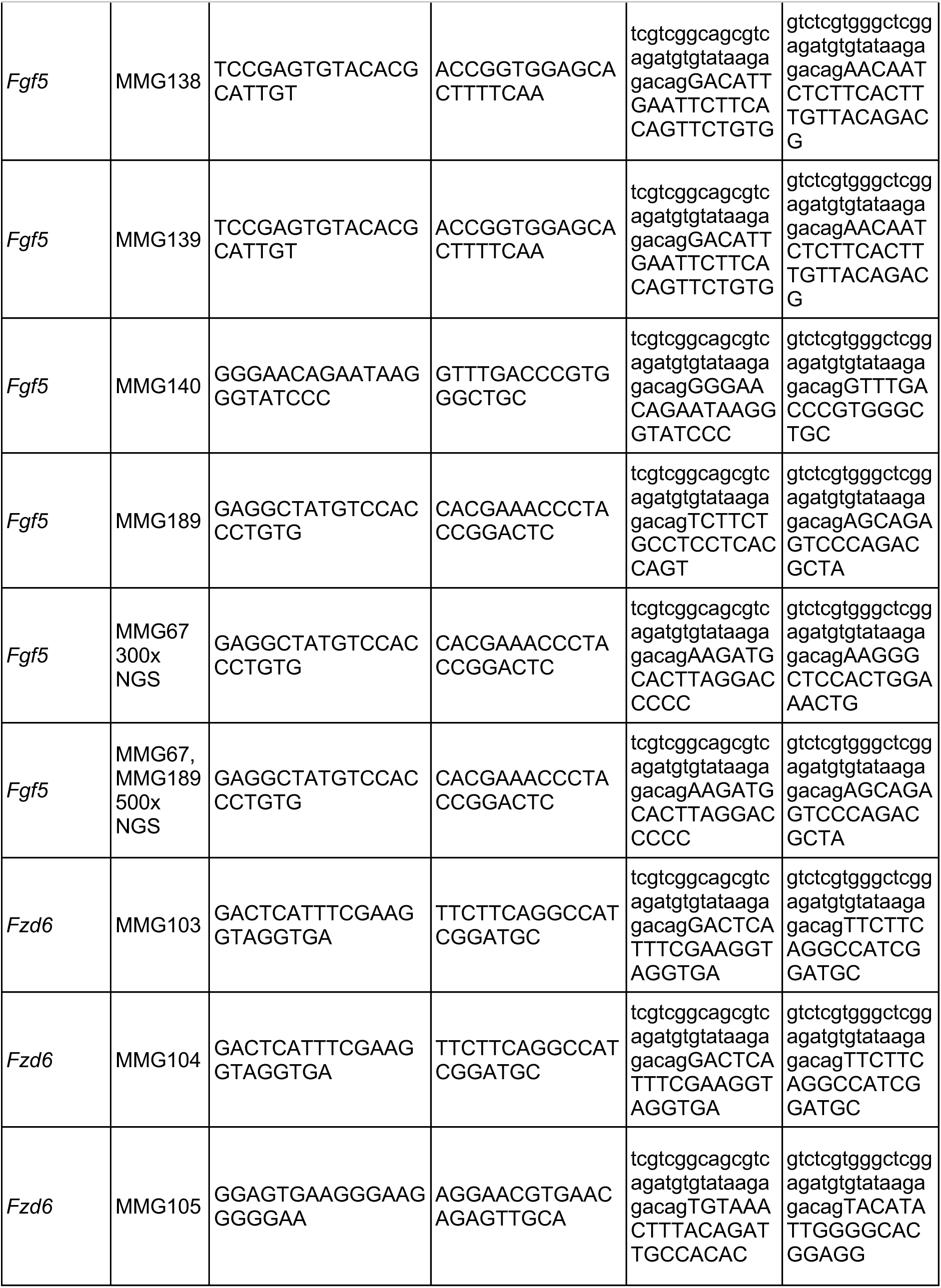
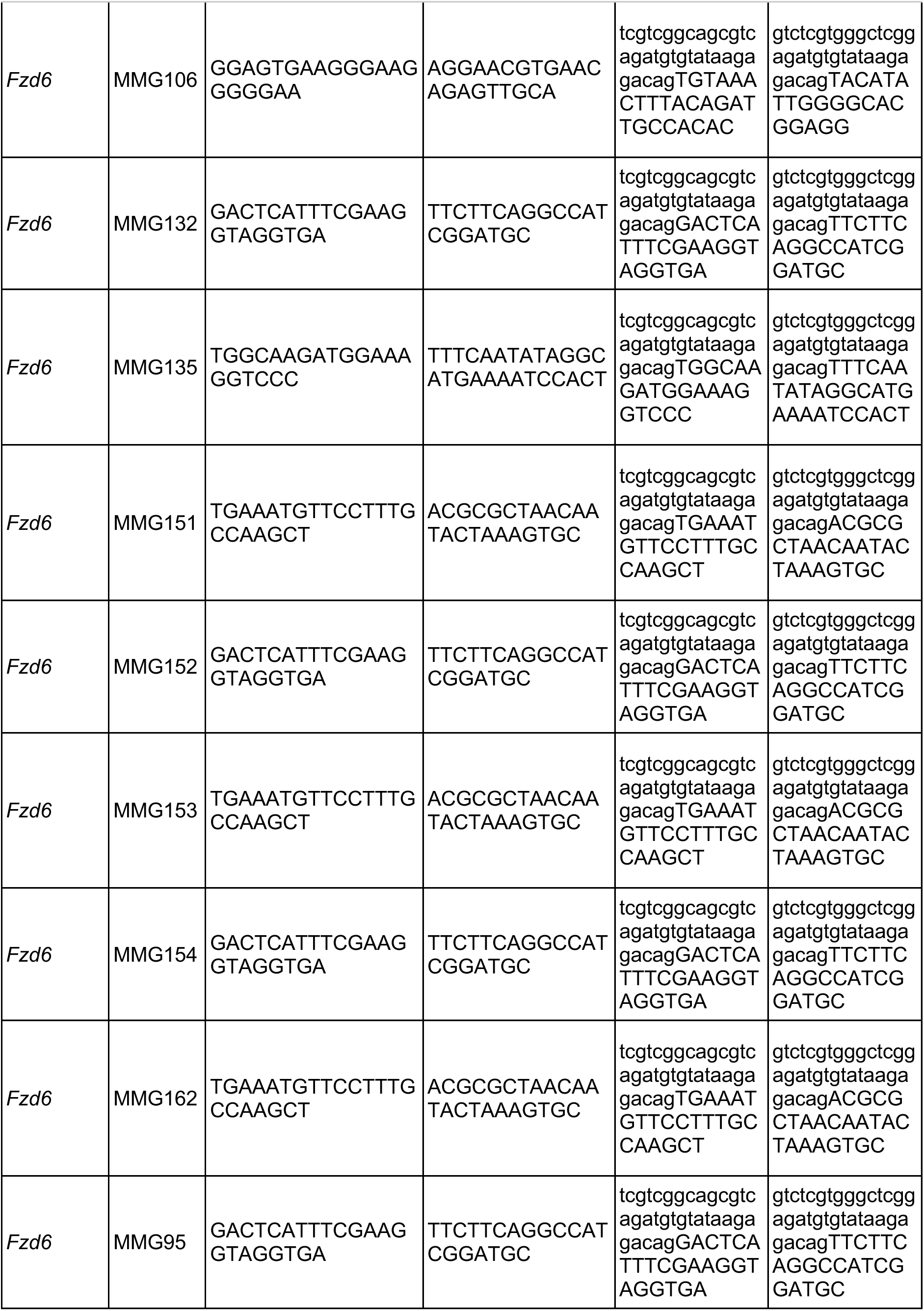
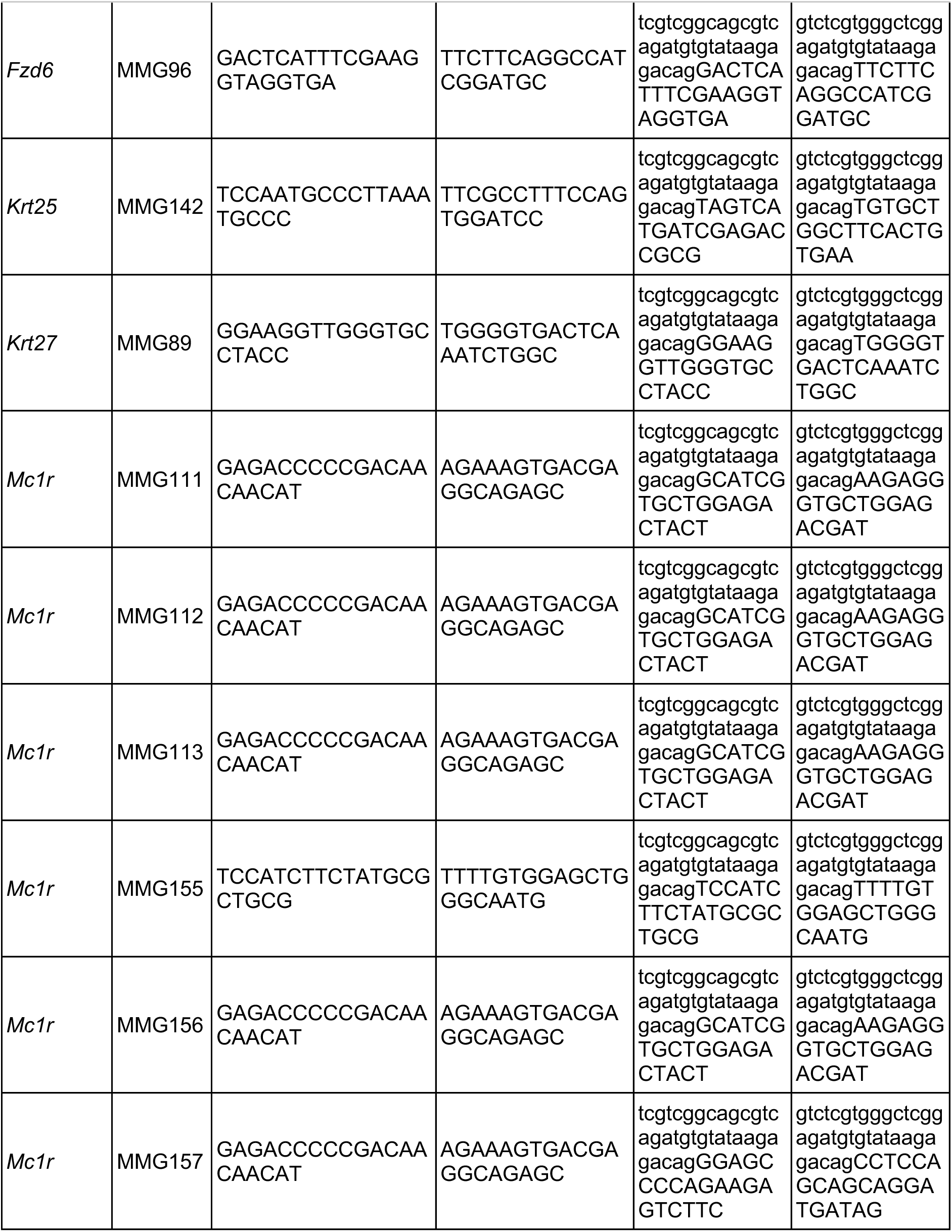
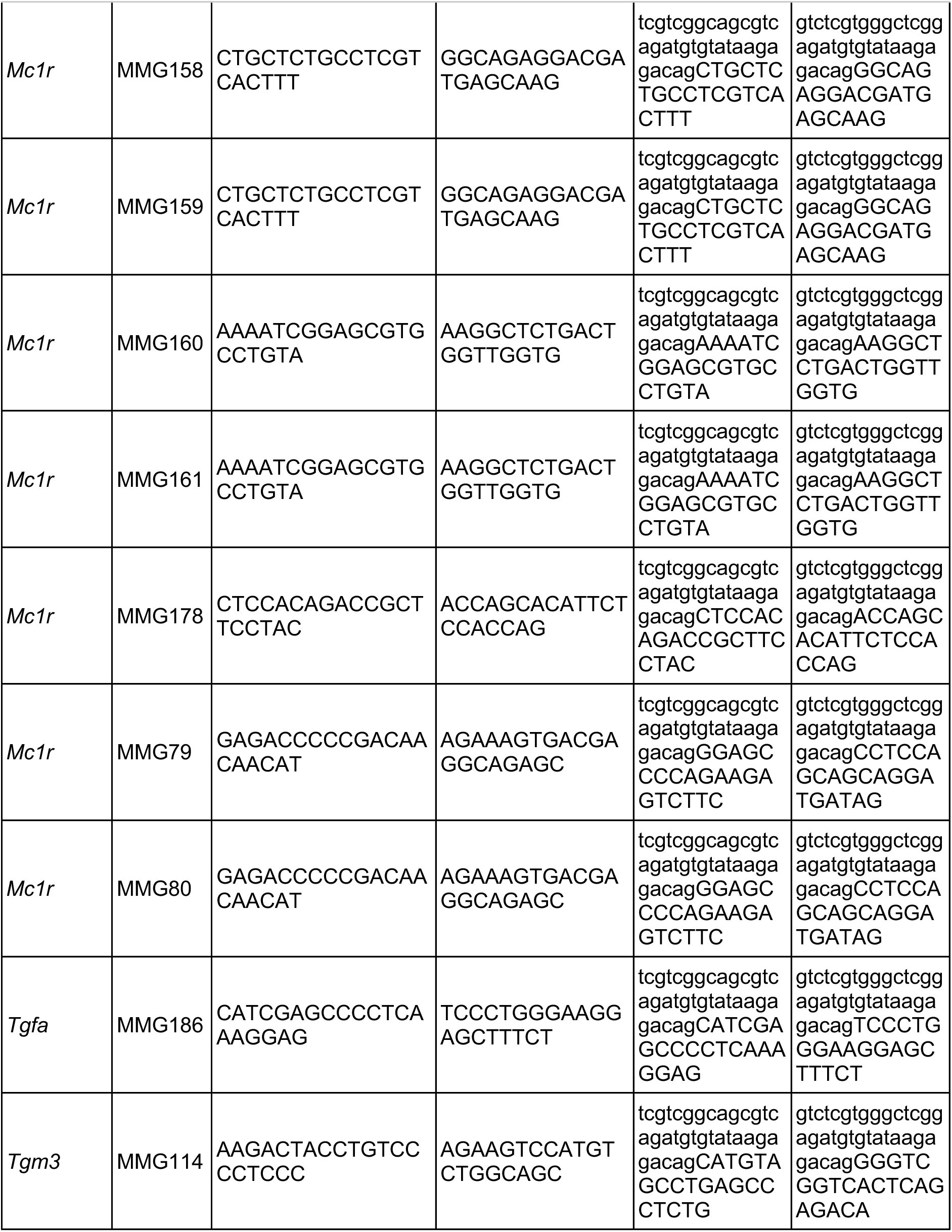
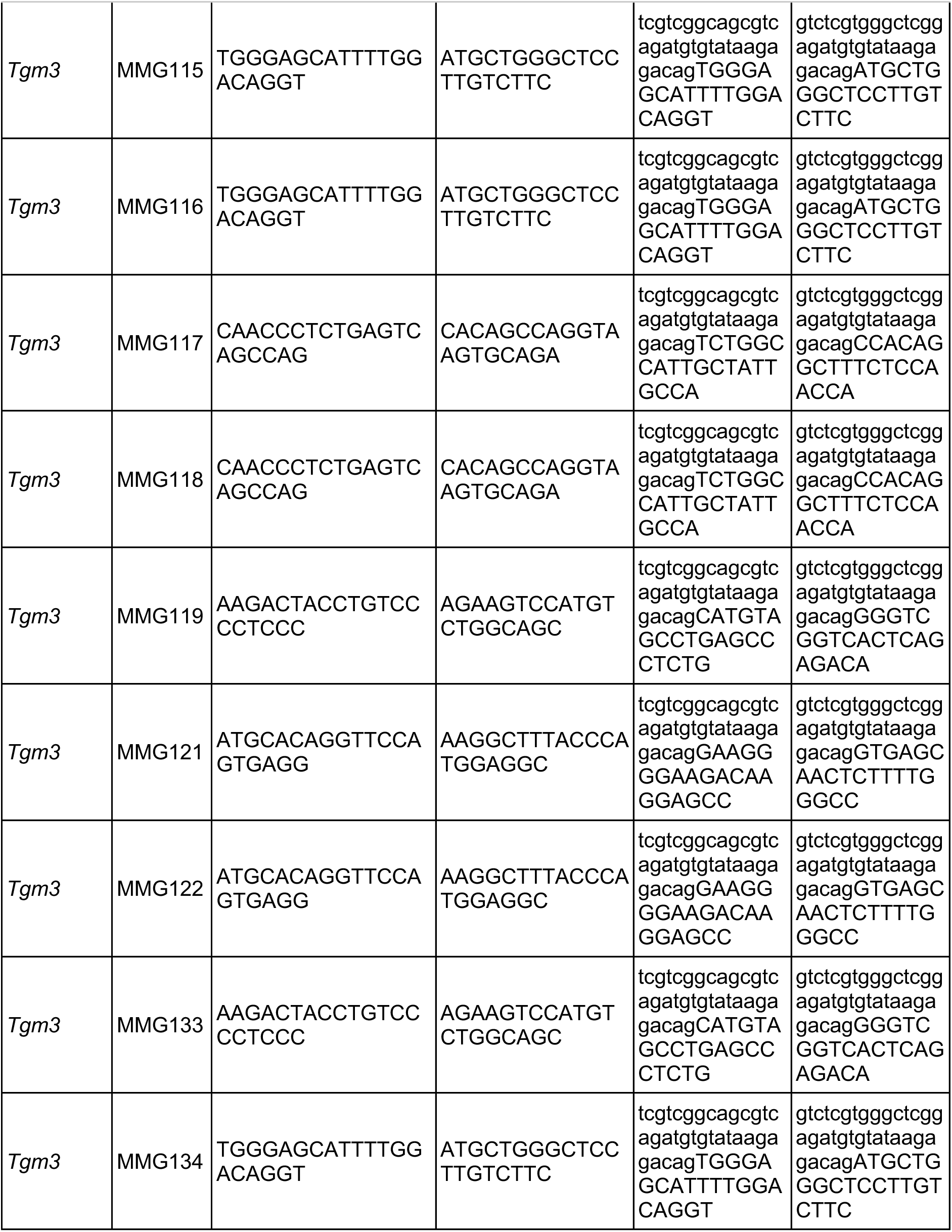
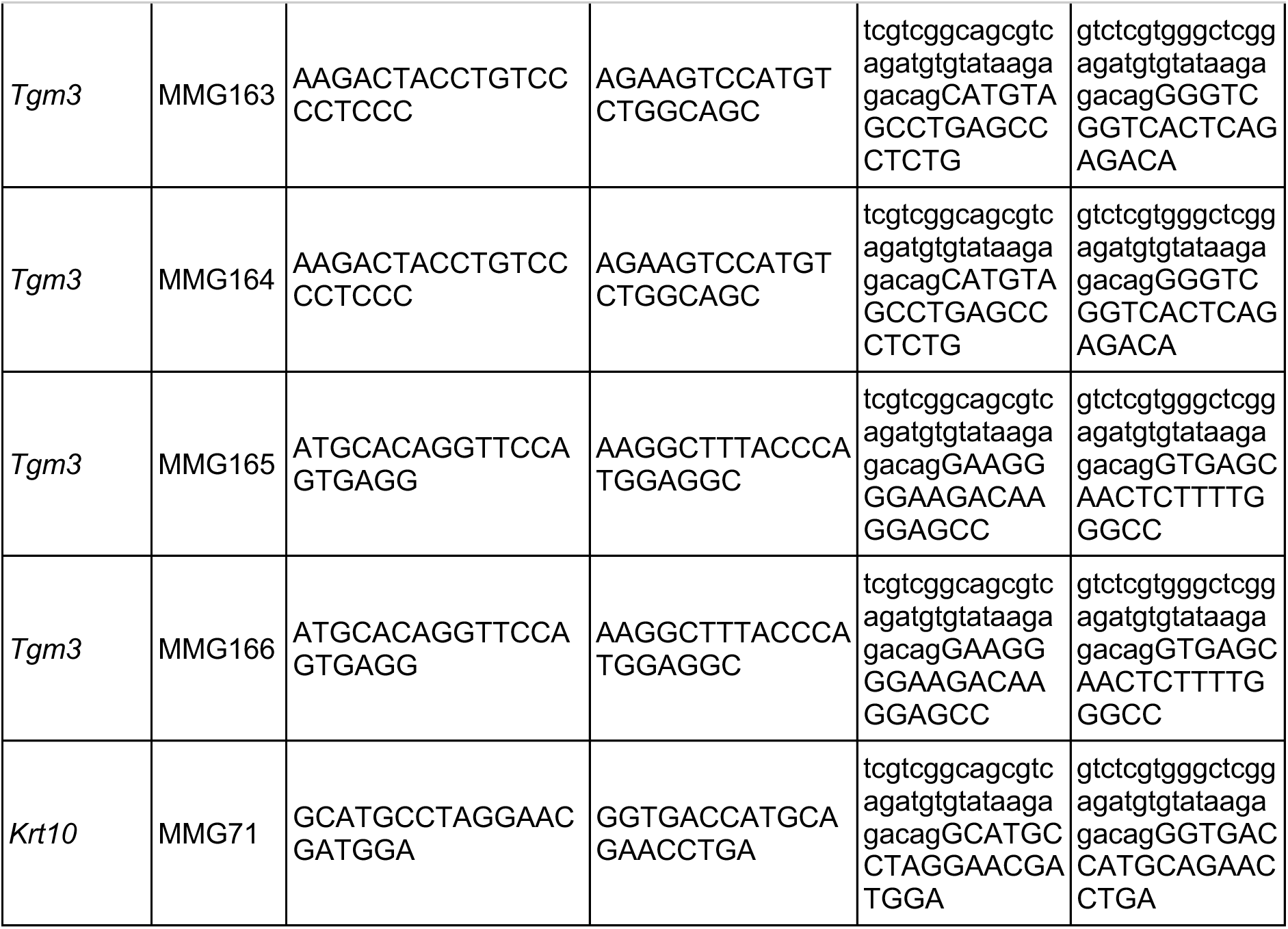
Genotyping primers for NGS and Sanger sequencing, related to Methods.

